# Temporal resolution of global gene expression and DNA methylation changes in the final phases of reprogramming towards induced pluripotency

**DOI:** 10.1101/547646

**Authors:** Michela Bartoccetti, Xinlong Luo, Ben van der Veer, Rita Khoueiry, Adrian Janiszewski, Jiayi Xu, Catherine Verfaillie, Vincent Pasque, Bernard Thienpont, Kian Peng Koh

## Abstract

The generation of induced pluripotent stem cells (iPSCs) involves activation of the endogenous pluripotency circuitry and global DNA demethylation late in reprogramming, but temporal resolution of these events using existing markers is insufficient. Here, we generated murine transgenic lines harboring dual fluorescent reporters reflecting cell-state specific expression of the master pluripotency factor *Oct4* and the 5-methylcytosine dioxygenase *Tet1*. By assessing reprogramming intermediates based on dual reporter patterns, we identified a sequential order of *Tet1* and *Oct4* gene activation at proximal and distal regulatory elements following pluripotency entry. Full induction of *Tet1* marks a pivotal late intermediate stage occurring after a phase of global gene repression, and preceding full activation of *Oct4* along with late naive pluripotency and germline-specific genes. Sequential activation of *Tet1* further distinguishes two waves of global DNA demethylation, targeting distinct genomic features and largely uncoupled from transcriptional changes, with dynamics unique to iPSC reprogramming. Moreover, we demonstrate that loss of *Tet1* is compatible with reprogramming towards full *Oct4* gene activation, but generates iPSCs with aberrant DNA methylation, chromosomal instability during lineage priming and defective differentiation potential. Therefore, the transcriptional logic of *Tet1* expression signals a deterministic epigenetic roadmap towards generation of high quality iPSCs.

## Introduction

Somatic cells can be reprogrammed *in vitro* into induced pluripotent stem cells (iPSCs) by enforced expression of transcription factors including the master pluripotency regulator OCT3/4 (also known as POU5f1 or OCT4)^1^. Pluripotent stem cells are able to self-renew indefinitely and to differentiate into any cell types of the body, and therefore have broad applications in fundamental research, disease modeling and cell-based therapies. During the generation of iPSCs, the de-differentiation of the somatic cell program to one closely resembling that of embryonic stem cells (ESCs) involves a dramatic remodeling of the epigenome in a highly inefficient process hampered by “epigenetic roadblocks” ^2^. In both murine and human reprogramming systems, activation of the *Oct4-*GFP reporter is often used as a gold standard for attaining “naive” pluripotency, a state of developmental potency equivalent with that of the inner cell mass (ICM) of the pre-implantation blastocyst. Nonetheless, the quality of even *Oct4-*GFP positive iPSCs can be highly variable, in part because of aberrant DNA methylation acquired during the reprogramming process ^3,4^. When present within gene promoters, DNA methylation is usually associated with long-term, stable gene repression and provides fundamental control of cell fate commitment ^5^. A better understanding of DNA methylation resetting during reprogramming is not only relevant to improving the quality of iPSC derivation, but also in understanding diseased states.

In recent studies to chart the molecular events of somatic cell reprogramming into iPSCs, reprogramming intermediates were isolated and characterized by using combinations of either cell surface markers, fluorescent reporters or single cell analysis ^6–11^. While THY1, SSEA1 and C-KIT offer reliable early- and intermediate- stage cell surface markers to isolate cell populations along the pathway of successful reprogramming ^8,11^, *Oct4-*GFP and *Nanog*-GFP reporters are the only commonly used markers for acquisition of pluripotency in murine systems ^6,12,13^. Nonetheless, dynamic changes in the transcriptome and chromatin, including global loss of DNA methylation during re-activation of the pluripotency network, may persist after entry into the pluripotent state ^8,14^. While much has been done to understand the early changes to erase somatic cell identity ^11,15,16^, relatively little is known about how global DNA methylation remodeling occurs in relation to gene expression to shape the final pluripotent cell identity late in the process. The transitional events in these late stages are especially difficult to resolve using existing markers, because only a small fraction of cells can be successfully reprogrammed to iPSCs among a heterogeneous bulk population that fails to.

In mammalian development, pluripotency is a feature of embryonic cells progressing in a continuum between the naive state of the pre-implantation ICM and the “primed” state of the gastrulation-poised post-implantation epiblast ^17^. From the latter, an alternative pluripotent cell type, known as epiblast stem cells (EpiSCs), can be directly derived in the mouse with more limited developmental potency compared to ICM-derived ESCs ^18,19^. Although OCT4 is expressed in both naive and primed pluripotent states, murine *Pou5f1* is regulated by distinct cell state-specific *cis* regulatory elements. A distal enhancer (DE) drives naive-state *Oct4* expression in pre-implantation embryos and in primordial germ cells (PGCs) but is inactive in the primed epiblast, where instead a proximal enhancer (PE) drives expression ^20^. At some point during reprogramming, exogenous expression of reprogramming factors becomes dispensable once endogenous *Oct4* expression is reactivated with the pluripotency network, presumably involving sequential activation of its PE followed by DE.

DNA methylation erasure at both *Oct4* distal and proximal elements is intricately associated with successful reprogramming to iPSCs ^1^. The activation kinetics of the DNA demethylation machinery may help to disentangle and understand these temporal events. The Ten-Eleven-Translocation (TET) family proteins, including TET1, TET2 and TET3, are the only known mammalian enzymes fully sufficient at driving pathways of active DNA demethylation, by converting 5-methylcytosine (5mC) to 5-hydroxymethylcytosine (5hmC) and further oxidation products at CpG dinucleotides ^21–23^. Of the three TET genes, *Tet1* expression is barely detectable in mouse embryonic fibroblasts (MEFs), but robustly up-regulated in iPSCs concomitantly with endogenous *Oct4* late in reprogramming ^8,24^. Because *Tet1* is a target gene regulated by OCT4 ^24,25^, a positive feedback logic involving *Tet1* and *Oct4* may function to jump-start the pluripotency circuitry during reprogramming ^26^. However, the sequential patterns of endogenous *Oct4* and *Tet1* gene activation during reprogramming remain unknown.

During early embryogenesis, *Tet1* expression is driven by two state-specific promoter-enhancer regions, using a similar *cis*-regulatory logic as for *Oct4*, where a distal promoter region spanning 6 kb drives an alternative transcription start site (TSS) at exon 1b which is activated exclusively in pre-implantation embryos and naive ESCs ^25^. A proximal promoter-enhancer coupling at exon 1a sustains *Tet1* expression through primed pluripotency. Here, we generated MEFs harboring distinct dual-fluorescent transgenic reporters for state-specific expression of *Oct4* and *Tet1*. By using live cell imaging and flow cytometry coupled with genome-scale analyses, our studies revealed a distinct trajectory of epigenetic events marking cell state transitions from early acquisition of pluripotency to terminal establishment of clonal iPSCs. We further clarify the role of TET1 in the reprogramming process by examining iPSCs generated from cells fully deficient of TET1. Surprisingly, TET1 is dispensable for *Oct4* activation, but affects proper epigenetic remodeling and quality of iPSCs.

## Results

### Characterization of transgenic reporter strains

We generated two independent murine transgenic strains harboring mCherry reporters to distinguish developmental stage-specific expression patterns of *Tet1* ^25,27^ (Figure 1A). The first strain, named Tg(*Tet1-*mCherry)B, contains a randomly integrated mCherry-2A-puromycin resistance cassette driven by a minimal 6-kb distal promoter region of *Tet1* (previously denoted as genomic region B6 in ref 25). From mouse v6.5 ES cell lines transfected with a plasmid containing this transgene, we selected two clonal lines (clones 37 and 43) that were validated by Southern blot and targeted locus amplification sequencing ^28^ to contain an intact transgene copy integrated at a single locus (Figure S1A-E).

**Figure 1.**
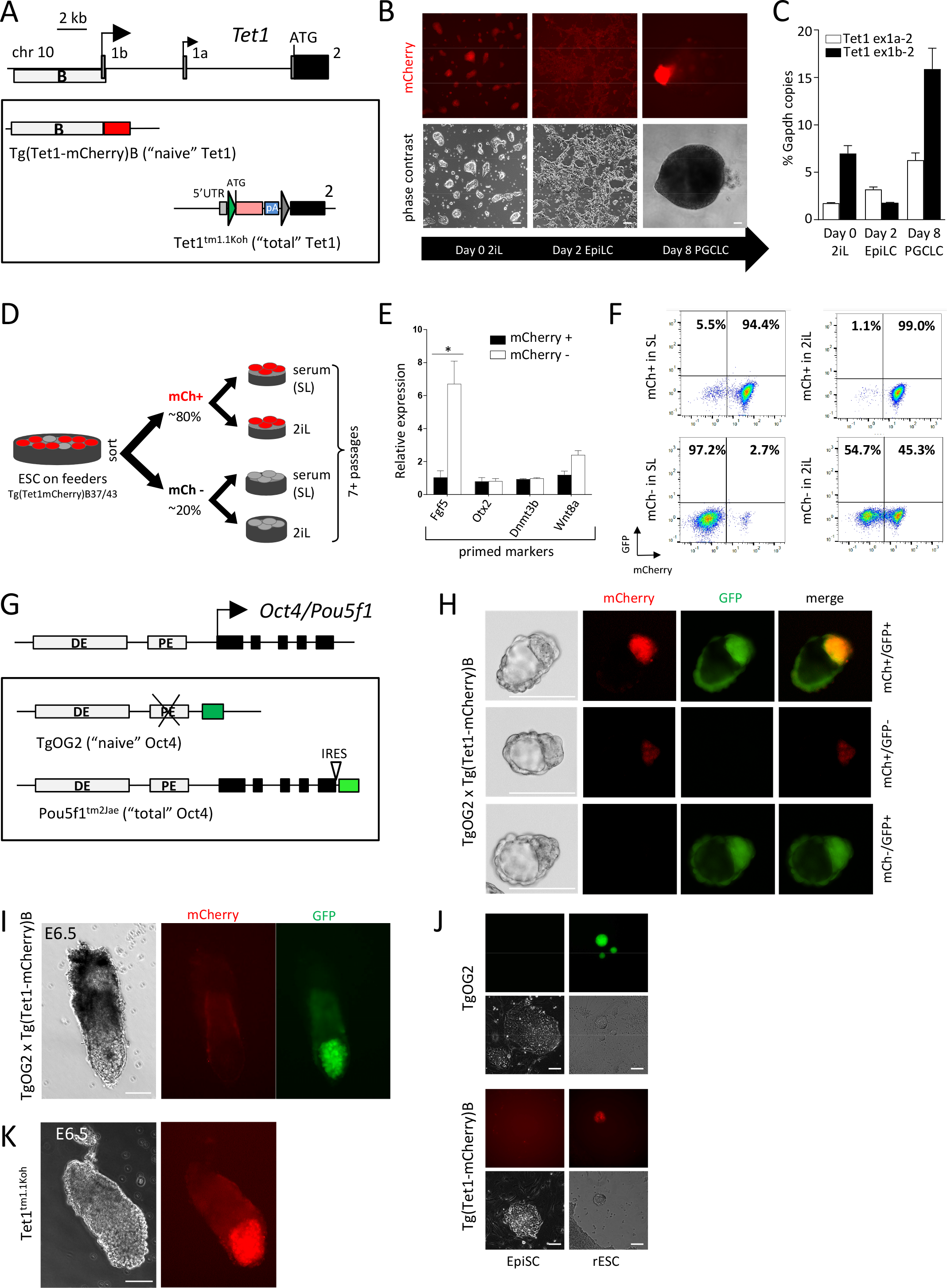
Characterization of transgenic reporter strains. (A) Schematic illustration of the 5′ genomic region of *Tet1*. 5’UTR and coding sequences are represented by grey and black boxes respectively. Bent arrows indicate alternative transcription start sites. The 6-kb distal promoter region B ^25^ is marked by a white rectangular bar. Below, schematic of *Tet1-*mCherry reporter lines. Red and pink boxes indicate the mCherry gene; green and grey triangles represent residual loxP and FRT sites respectively after recombination. (B) Fluorescence and phase contrast images of Tg(*Tet1-*mCherry)B cell lines during differentiation towards primordial germ cell-like cells (PGCLCs). (C) Expression of *Tet1* 5’ transcript isoforms (alternative exons 1a or 1b spliced to exon 2) during PGCLC differentiation. Quantitative PCR expression values are shown as mean ± SEM (n=3). (D) Schematic overview of cell sorting and re-culture in different media conditions to assess transgene activity in Tg(*Tet1-*mCherry)B lines. In serum and feeder cultures, ~20% of cells from both clonal lines are mCherry-negative by FACS analysis. (E) Relative expression of primed pluripotency marker genes in mCherry negative and positive populations. Expression values are means ± SEM (n=3; n=2 for clone 37 and n=1 for clone 43).* *P* < 0.0001. (F) Representative flow cytometry plots of sorted mCherry-negative and -positive cells after re-culture in serum and 2i conditions for more than 7 passages. (G) Schematic illustration of 5’ genomic region of *Oct4/Pou5f1*. Distal enhancer (DE) and proximal enhancer (PE) elements are indicated as white boxes, exons in black boxes. Below, schematic of *Oct4-*GFP reporter lines. Green boxes (in two shades) indicate GFP. (H-K) Fluorescence and bright field images. (H) Pre-implantation blastocysts from TgOG2 × Tg(*Tet1-*mCherry)B strain crosses. Top row shows an embryo with dual fluorescence and lower two rows are single reporter littermate embryos. (I) Post-implantation E6.5 TgOG2 × Tg(*Tet1-*mCherry)B embryos. (J) EpiSCs and “reset” rESCs in single-reporter TgOG2 and Tg(*Tet1-*mCherry)B lines. (K) E6.5 *Tet1*^tm1.1Koh^ embryos. All scale bars in Figure 1 represent 100 μm.

To characterize the Tg(*Tet1-*mCherry)B construct for naive-specific expression, we observed mCherry fluorescence of the transgenic reporter cell lines propagated in different pluripotency states. mCherry fluorescence was detectable in ESCs cultured under self-renewing conditions sustained by leukemia inhibitory factor (LIF) in either serum-containing media (SL) or defined media containing inhibitors of MEK and GSK (2iL); the latter maintain cells in a pluripotent ground state ^29^. Once cells in ground state were differentiated to primed epiblast-like cells (EpiLCs) upon supplementation with basic fibroblast growth factor (bFGF) and Activin A, mCherry fluorescence was rapidly downregulated in all cells (Figure 1B). Further differentiation into CD61+/SSEA1+ sorted primordial germ cell-like cells (PGCLCs) ^30^ expressing *Stella* and *Blimp1* (Figures S1F-G) reactivated mCherry expression (Figure 1B). In congruence with mCherry reporter expression, *Tet1* transcripts initiated from the alternative exon 1b decreased in EpiLCs and increased dramatically in PGCLCs (Figure 1C). In contrast, transcripts initiated from a constitutively, albeit weakly, active promoter at exon 1a ^25^ did not correlate with gain or loss of the naive state (Figure 1C).

Although Tg(*Tet1-*mCherry)B activation appeared restricted to naive cells, fluorescence-activated cell sorting (FACS) showed that the reporter was silenced in ~20% of ESCs cultured in serum and LIF (SL) in both clonal lines ^25^. To examine this mCherry-negative population, we sorted cells into mCherry-positive and negative fractions followed by re-culture in either SL or 2iL conditions (Figure 1D). Quantitative reverse transcription-PCR (Q-PCR) analysis of sorted fractions indicated that the mCherry-negative cells expressed steady-state endogenous exon 1b transcripts and naive pluripotent markers such as *Klf4* or *Esrrb* at levels equivalent to that observed in mCherry-positive cells (Figure S1H-I); however, the primed epiblast marker *Fgf5* was clearly up-regulated in mCherry-negative and not in mCherry-positive cells (Figure 1E). When mCherry-positive cells were re-cultured in SL or in 2iL conditions, they mostly retained mCherry expression. By contrast, mCherry-negative cells remained mostly negative in SL, but about 45% could re-activate mCherry reporter when re-cultured in 2iL (Figures 1F and S1J). Although these fluctuations in mCherry expression may indicate a degree of random transgene inactivation independent from the endogenous gene, our cell sorting analysis nonetheless suggests that the Tg(*Tet1-*mCherry)B reporter can distinguish subtle differences in transcriptionally heterogeneous pluripotent cell populations in metastable ESC serum cultures. Moreover, any transgene silencing in the presence of serum is reversible in 2iL conditions promoting the naive pluripotent ground state.

Because both clonal Tg(*Tet1-*mCherry)B cell lines 37 and 43 reported naive state similarly *in vitro*, we proceeded with blastocyst injections of both ESC clones to generate two independent B6;129S-Tg(*Tet1-*mCherry)B mouse strains. To determine how the naive-specific expression pattern of the transgene -named “naive” *Tet1-*mCherry - relates to that of *Oct4*, we compared the “naive” *Tet1-*mCherry strains with B6;CBA-Tg(*Pou5f1*-EGFP)2Mnn/J (or TgOG2), a commonly used *Oct4-*GFP reporter for naive pluripotency ^31^. In TgOG2 mice, a randomly integrated 18-kb transgene contains the *Oct4* genomic locus, wherein PE is deleted, driving expression of an enhanced green fluorescent protein (EGFP) gene (Figure 1G). This strain - named “naive” *Oct4-*GFP - is distinct from an independent *Oct4-*GFP strain (B6;129S4-*Pou5f1*^tm2Jae/J^) which harbors a gene-targeted knock-in of EGFP downstream of an internal ribosomal entry site (IRES) between the stop codon and endogenous polyA signal ^32^ (Figure 1G). Because the *Oct4-*IRES-GFP modification reflects gene activity by all upstream *cis* regulatory elements, we refer to this strain as “total” *Oct4-*GFP.

In either single or dual reporter mouse embryos obtained from strain inter-crossings of Tg(*Tet1-*mCherry)B and TgOG2, we detected “naive” *Oct4-*GFP and “naive” *Tet1-*mCherry signals in the ICM of pre-implantation stage blastocysts (Figure 1H). In post-implantation E6.5 embryos, “naive” *Tet1-*mCherry expression was no longer detectable as expected. However, “naive” *Oct4-*GFP was still detectable at E6.5 and persisted until E7.5 (Figures 1I and S1K). This persistence of *Oct4-*GFP in post-implantation TgOG2 embryos, in contrast to the earlier reported pre-implantation stage-specific expression of the original *Oct4-*lacZ transgene ^20^, has also been observed by others ^33^ and may be a consequence of the long half-life of EGFP *in vivo*. Nonetheless, both naive *Tet1* and *Oct4* reporters were silenced in EpiSCs derived from E6.5 embryos and maintained in the presence of Activin A and bFGF, indicating that *in vitro* cultures of the primed state effectively turn off both naive reporters (Figure 1J). Using a protocol to convert EpiSCs to “reset-naive” ESCs (rESC) *in vitro* by overexpression of Nr5a1^34^, we observed that both “naive” *Tet1-*mCherry and *Oct4-*GFP reporters could be reactivated upon reversion to naive-like states, also evident as reacquisition of strong alkaline phosphatase activity (Figures 1J and S1L). Collectively, both *in vitro* and *in vivo* conditions confirmed the specificity of the “naive” *Tet1-*mCherry reporter for the naive pluripotent state.

To mark *Tet1* gene activity from both proximal and distal promoter regions, we generated a second independent *Tet1-*mCherry strain, the B6-*Tet1*^tm1.1Koh^ line, in which the mCherry reporter was targeted after the ATG start codon of the coding sequence in exon 2 (Figure 1A). This strain – named “total” *Tet1-*mCherry - was generated by reporter exchange in the original *Tet1-*loxP-lacZ-loxP-mCherry knock-in cassette of the *Tet1*^tm1Koh^ allele ^27^ following breeding with Cre-deleter mice to obtain a *Tet1-*loxP-mCherry allele. In agreement with *Tet1* expression in the post-implantation epiblast ^25,27^, transgenic embryos showed detectable signals in the E6.5 epiblast (Figure 1K).

### Kinetics of *Tet1* and *Oct4* gene activation during reprogramming

To examine dual reporter patterns during iPSC reprogramming, we bred dual transgenic strains that harbor distinct combinations of *Tet1-*mCherry and *Oct4-*GFP reporters to isolate MEFs as the starting somatic cells. First, we crossed the “naive” *Tet1-*mCherry strain with “naive” *Oct4-*GFP (TgOG2) to obtain MEFs containing a “naive × naive” reporter combination (Figure 2A). Fluorescence from both reporters was not detectable in transgenic MEFs. Upon lentiviral transduction of an inducible *Oct4-*F2A-*Klf4*-IRES-*Sox2*-E2A-*cMyc* (OKSM) cassette ^35^ and a reverse tetracycline-controlled transactivator (rtTA), we monitored the time-course of reprogramming at daily intervals by flow cytometry. We first detected mCherry+/GFP− cells from day 10 and progressively more mCherry+/GFP+ cells until the last time-point at day 17 (Figure 2B and 2C).

**Figure 2.**
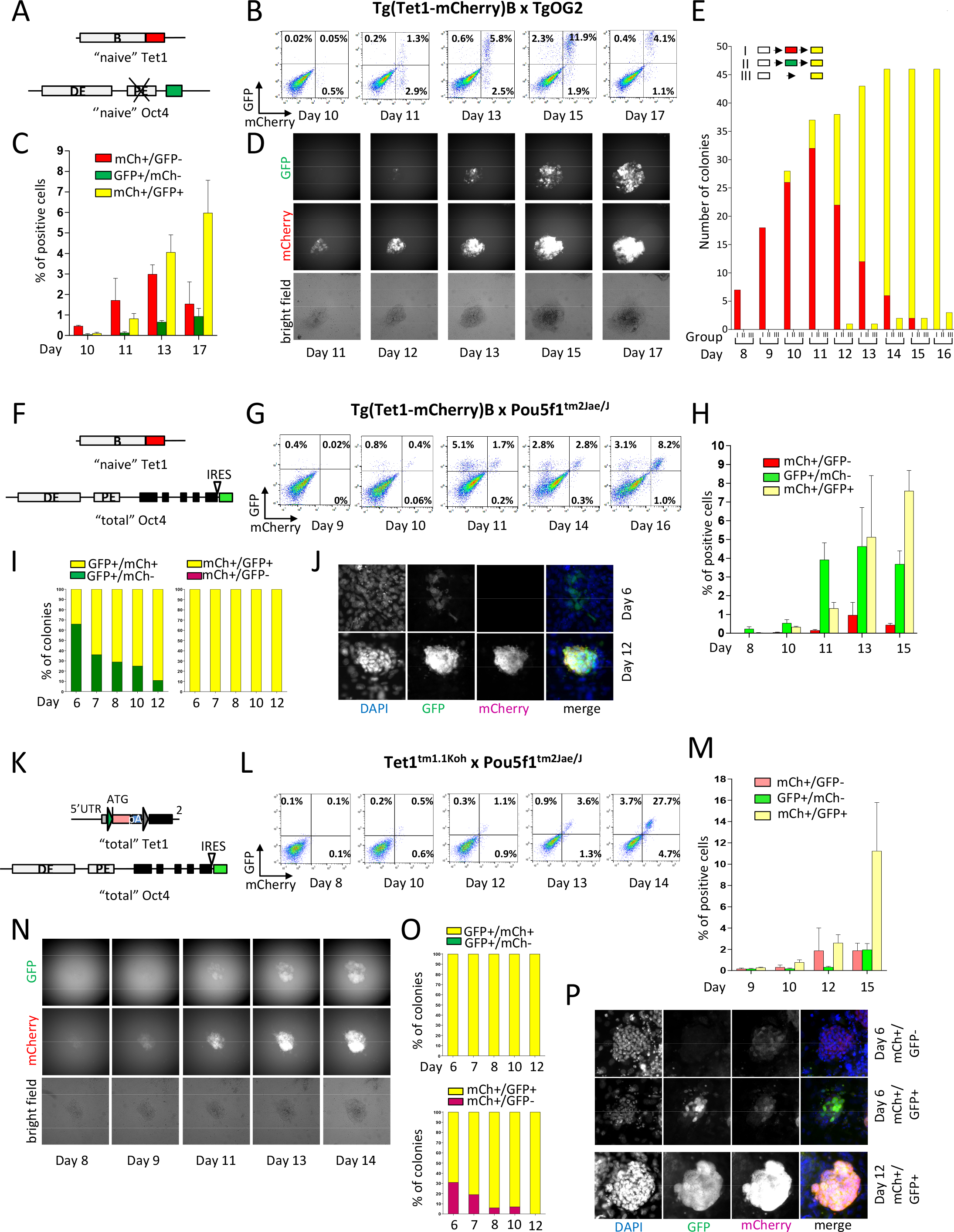
Kinetics of *Tet1* and *Oct4* gene activation during reprogramming. (A) Schematic of dual transgenic Tg(*Tet1-*mCherry)B; Tg(*Pou5f1*-EGFP)Mnn line. (B-C) Flow cytometry of cells harvested at different time points during reprogramming. Percentages of gated cell fractions are shown in C as mean ± SEM (n=3 independent assays). (D) Representative fluorescence images of a single colony imaged over time. See Figure S2A for stitched images of multiple colonies. (E) Colony counts based on sequential activation patterns of fluorescent-reporters. Colonies were assigned to 3 groups based on whether they activate mCherry (Group I) or GFP (Group II) first, before turning double positive, or concurrently activate both on the same day (Group III). See Figure S2B for scoring in a replicate experiment. (F-H) Same as A-C for Tg(*Tet1-*mCherry)B; *Pou5f1*^tm2Jae/J^ line. (I-J) Percentage of colonies (I) and immunofluorescence images (J) for GFP and mCherry co-expression during reprogramming of Tg(*Tet1-*mCherry)B; *Pou5f1*^tm2Jae/J^ line. (K-N) Same as A-D for *Tet1*^tm1.1Koh^; *Pou5f1*^tm2Jae/J^ line. See Figure S2E for stitched images. (O-P) Percentage of colonies (O) and immunofluorescence images (P) for GFP and mCherry co-expression during reprogramming of *Tet1*^tm1.1Koh^; *Pou5f1*^tm2Jae/J^ line.

To track the kinetics of *Tet1* and *Oct4* reactivation at the single colony level, we performed live cell imaging, acquiring images of reprogramming cultures in defined fields from day 7 to day 17 on a daily basis. Based on all colonies imaged on day 16-17, we retrospectively tracked images of each colony until day 8 and scored the appearance of mCherry and GFP signals per day (Figure 2D and S2A). Because ~98% of colonies showed strong double-positive mCherry+/GFP+ signals by day 16, we classified scored colonies into three predicted groups: (I) *Tet1-*mCherry detectable before *Oct4-*GFP; (II) *Oct4-*GFP activation preceded *Tet1-*mCherry and (III) both *Oct4-*GFP and *Tet1-*mCherry were co-activated on the same day.

In agreement with the FACS data, the majority of the colonies activated “naive” *Tet1-* mCherry first, followed by *Oct4-*GFP (red turning into yellow bars, Figures 2E and S2B). These sequential events were separated temporally by at least a one-day gap (Group I) (Figure 2D-E and S2A-B). There were no detectable colonies that activated “naive” *Oct4-*GFP followed by “naive” *Tet1-*mCherry (Group II); a few (< 10%) co-activated the two reporters on the same day (Group III).

In very rare cases (~2% of colonies of >100 scored), we observed mCherry−/GFP+ colonies that failed to reactivate *Tet1-*mCherry even upon colony isolation and expansion. On the other hand, we did not detect any colony that activated *Tet1-*mCherry but failed to activate *Oct4-* GFP. Q-PCR analysis of two mCherry−/GFP+ clonally expanded iPSC lines demonstrated that endogenous transcripts from exon 1b could be detected at comparable levels as in double-positive clonal lines, in line with transgene silencing of *Tet1-*mCherry occurring at low frequencies in reprogramming cells. These mCherry−/GFP+ lines were not examined further because they rarely occur in our reprogramming assays, but may possibly represent iPSCs of poorer grade.

Scoring of individual colonies by live cell imaging preserved information at the clonal level and with a temporal history. The overall picture convincingly showed that the vast majority of cells that successfully form iPSC colonies follow a distinct trajectory of activating naive-specific *Tet1* expression prior to naive-specific *Oct4* expression.

Next, we inter-crossed the “naive” *Tet1-*mCherry and “total” *Oct4-*GFP strains to obtain “naive × total” dual transgenic MEFs (Figure 2F). By flow cytometry, we detected mCherry−/GFP+ populations appearing first around day 9-10 and only a minor fraction of mCherry+/GFP− cells after day 11, whereas mCherry+/GFP+ cells progressively accumulated from day 10 until day 16 (Figure 2G and 2H). In time-course observations by live fluorescence imaging, the majority of colonies (45.9%) appeared to gain GFP and mCherry signals simultaneously (Group III) (Figure S2C-D). The fractions of colonies that appeared to activate *Oct4-*GFP prior to *Tet1-*mCherry (Group II) exceeded those that activated mCherry before GFP (Group I) (Figure S2D). Similar results were obtained using both clonal strains from Tg(*Tet1-*mCherry)B37 and B43 of both genders, and at different generations of backcrossing (N2-N4) or in genetically heterogeneous CD1 background (Figure S2D).

To rule out that these observations were skewed by the higher background fluorescence of the *Oct4-*IRES-GFP reporter, we used GFP and mCherry antibodies to perform immunofluorescence (IF) detection in reprogramming cells fixed at different time-points to validate our findings. To facilitate reprogramming of cells seeded on coated coverglass in this assay, we added ascorbic acid to the serum-containing media to enhance reprogramming efficiency ^36^. By this independent assay, we identified GFP-positive colonies and scored whether these stained for mCherry, and vice versa in mCherry-positive colonies, on the premise that the earlier marker should appear in colonies that are negative for the other marker. Indeed, >60% of cell clusters appearing early at day 6 were GFP+/mCherry−; as reprogramming progressed until day 12 (the last assayed time-point), the proportion of GFP+/mCherry+ colonies increased (Figure 2I, left and 2J). In contrast, mCherry+ colonies were always GFP+ (Figure 2I, right). These results suggest that the first evidence of endogenous *Oct4* gene activity occurs slightly before the activation of the naive-specific *Tet1* distal promoter.

To find out the temporal order of “total” *Tet1* gene activation relative to “total” *Oct4* activation, we used a third combination of dual reporters obtained from strain intercrosses of B6-*Tet1*^tm1.1Koh^ and B6;129S4-*Pou5f1*^tm2Jae/J^ (Figure 2K). FACS analysis showed a parallel activation of mCherry and GFP around day 12 with a minor fraction gated as mCherry+/GFP− (Figure 2L-M). In live colony images, we detected earlier appearance of mCherry signals than GFP in almost all colonies scored (Figure 2N and S2E). IF assays further verified detection of a fraction (~30% at day 6) of mCherry-positive colonies that were GFP-negative; on the other hand, all GFP-positive colonies are doubly positive for mCherry (Figure 2O-P), suggesting that total *Tet1* gene activity is detectable slightly before that of “total” *Oct4*.

While temporal differences in reporter activation may be caused by strain differences in the starting cells, we ruled that out as a major confounding factor by comparing GFP detection in both “naive” Tet1 × “total” Oct4 and “total” Tet1 × “total” Oct4 strains, both harboring the *Oct4-*IRES-GFP reporter, during reprogramming. In parallel IF assays performed during a reprogramming time-course, both strains showed GFP signals detectable in similar numbers of colonies from day 6 (Figure S2F), suggesting similar activation kinetics of the *Oct4-*IRES-GFP allele when bred to either mixed B6;129 or incipient congenic B6 backgrounds. Collectively, our observations suggest that both *Tet1* and *Oct4* loci are activated at proximal regulatory elements nearly synchronously during reprogramming, with *Tet1* marginally earlier than *Oct4*, followed by a second phase of activation of distal regulatory domains, in which naive-specific *Tet1* gene activity more clearly preceded that of *Oct4*.

### Activation dynamics in different reprogramming conditions and relative to NANOG and DPPA4

Since reprogramming variability may potentially be attributable to multiple copies of lentiviral OKSM integration, we further tested our dual reporters in an OKSM reprogrammable mouse system [B6;129S4-Col1a1^tm1(tetO-*Pou5f1*,-*Klf4*,-*Sox2*,-*Myc*)Hoch/J^], which harbors the inducible 4-factors integrated as single copy in every cell ^37^. By crossing dual reporter strains to OKSM mice, we generated triple heterozygous MEF lines for reprogramming assays following lentiviral infection of M2-rtTA and doxycycline treatment ^38^. Because reprogramming efficiencies can be boosted by the use of defined media containing knockout serum replacement (KSR) and fresh supplementation of ascorbic acid ^36,39^, we compared reporter activation kinetics in three different LIF-containing ES media compositions: (i) serum, (ii) serum supplemented with ascorbic acid (serum+AA) and (iii) KSR supplemented with ascorbic acid (KSR+AA).

We focused on the “naive × naive” (i.e. “naivex2”) and “total × total” (i.e.”totalx2”) combinations, to verify if every iPSC colony scored progressed from activating *Tet1-*mCherry followed by *Oct4-*GFP. In both dual reporter combinations, final iPSC colony counts were increased by the addition of ascorbic acid to serum-containing media, and boosted further when serum was replaced by KSR (Figure S3A-D). Although the kinetics of the fluorescence activation was faster in KSR+AA, the sequential order remained the same: *Tet1-*mCherry was reactivated first, followed by *Oct4-*GFP in both dual reporter systems (Figure 3A-B). Especially in the OKSM;Tg(*Tet1-*mCherry)B;TgOG2 (“naivex2”) strain, the temporal gap between the reactivation of mCherry and GFP was distinct in every colony when compared to cells reprogrammed by the lentiviral system (Figure S3E compared to Figure 2D), such that Group III colonies were no longer detected.

**Figure 3.**
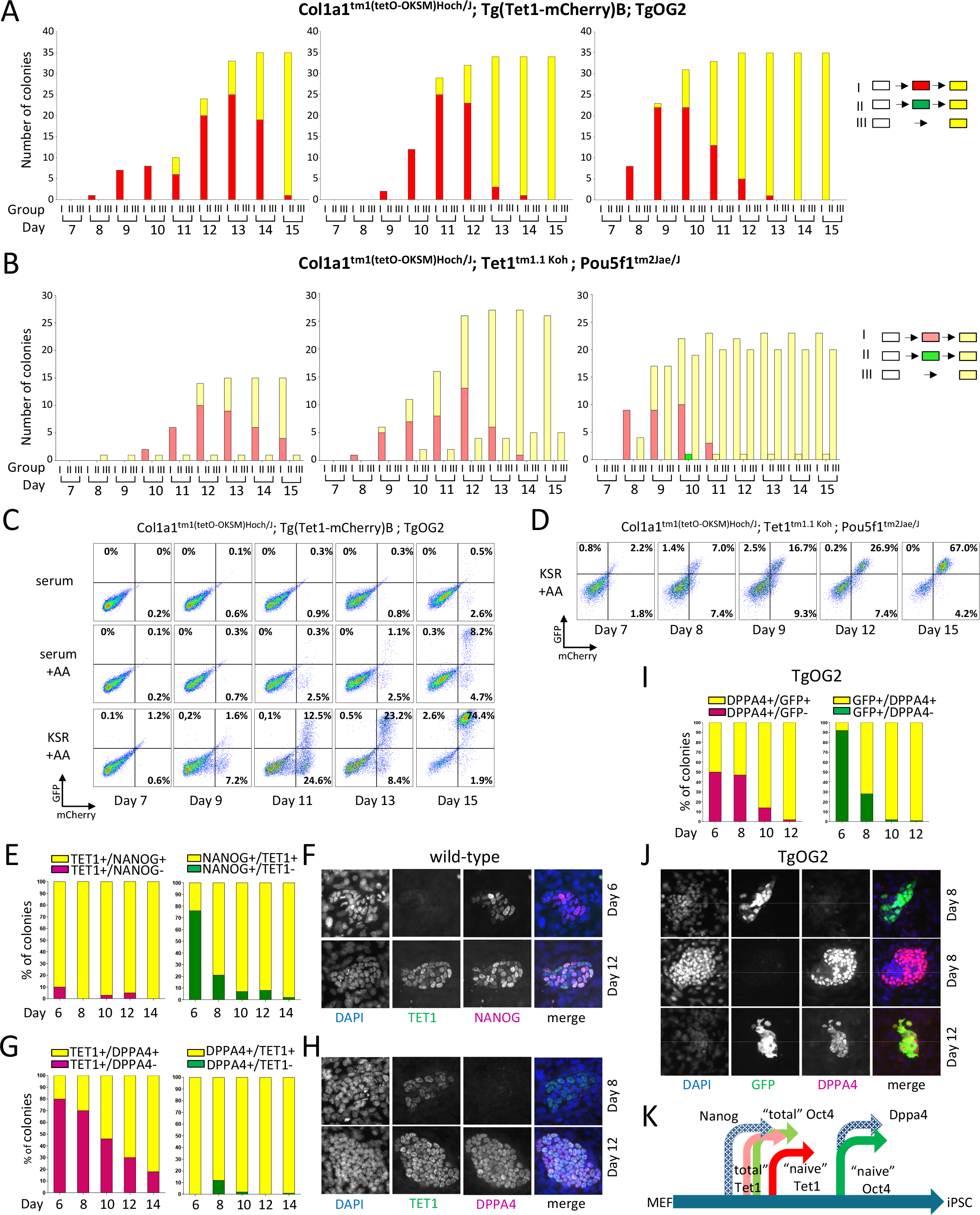
Activation dynamics in different reprogramming conditions and relative to NANOG and DPPA4. (A-B) Daily colony counting based on fluorescence activation in different media conditions of Tg(*Tet1-*mCherry)B; Tg(*Pou5f1*-EGFP)Mnn (“naivex2”) (A) and *Tet1*^tm1.1Koh^; *Pou5f1*^tm2Jae/J^ (“totalx2”) dual reporter lines (B). (C-D) FACS analysis of cells harvested at different time points in serum, serum + ascorbic acid (AA) and knockout serum replacement (KSR) +AA for Tg(*Tet1-*mCherry)B; Tg(*Pou5f1*-EGFP)Mnn (C) and in KSR+AA for *Tet1*^tm1.1Koh^ × *Pou5f1*^tm2Jae/J^ (D). (E-J) Percentage of colony counts and immunofluorescence images for TET1 and NANOG (E-F), TET1 and DPPA4 (G-H) and GFP and DPPA4 (I-J) co-expression during reprogramming. (K) Schematic overview of the temporal order of pluripotency gene activation.

Next, we used flow cytometry to profile dual-reporter fluorescence patterns of reprogramming cell populations in the three media conditions on a daily basis. With single-copy/allele OKSM expression, reprogramming efficiency was very low in serum-containing media, even in “naivex2” cells, in which only 0.5% of cells were double positive at day 15. While serum+AA increased that population to 8.2%, efficiency was greatly boosted to 74.4% in KSR+AA, in which a distinct cluster of mCherry+/GFP− population appeared as early as day 9 (Figure 3C). Similarly, the efficiency of OKSM; *Tet1*^tm1.1Koh^; *Pou5f1*^tm2Jae/J^ dual reporter cells (“totalx2”) acquiring double mCherry+/GFP+ fluorescence by day 15 in KSR+AA reached 67% of the total population. Notably, the single-fluorescent mCherry+/GFP− fraction was discernable as early as day 8-9 (Figure 3D and S3F).

Among pluripotency factors, *Nanog* is known to be expressed early, while the naive marker *Dppa4* is activated late in reprogramming ^10,13^. To assess *Tet1* expression relative to *Nanog*, we stained fixed cells at different time points during reprogramming in serum+AA using antibodies for NANOG and TET1 ^40^. Colonies positive for TET1 were almost all already positive for NANOG. However, a majority (~70%) of NANOG-positive cell clusters detectable at the early day 6 time-point showed no detectable staining for TET1 (Figure 3E-F). These observations suggest that NANOG is expressed earlier than TET1, although the temporal gap is likely to be narrow. Next, we co-stained for DPPA4 and TET1. In this case, TET1-positive colonies were mixes of DPPA4-positive and negative, with more DPPA4-negative colonies between day 6-10, and progressively more double-positive colonies by days 12-14 (Figure 3G-H). On the other hand, DPPA4-positive colonies were almost all TET1-positive. These profiles clearly suggest that DPPA4 is detectable after TET1 accumulates.

To further distinguish naive state-specific versus total *Tet1* expression, we co-stained cells reprogrammed from Tg(*Tet1-*mCherry)B (“naive Tet1”) MEFs for mCherry or total TET1. As expected, weak total TET1 expression was detected in days 6 and 8 colonies, clearly preceding mCherry expression that accumulated later (Figure S3G-H). In co-IF staining for DPPA4 and mCherry, DPPA4 expression was detectable in colonies after appearance of mCherry (Figure S3I-J). Finally, we performed co-IF staining for DPPA4 and GFP during reprogramming of TgOG2 (“naive Oct4”) MEFs to determine which is the last naive marker activated. We detected a mix of early intermediate colonies that either co-expressed DPPA4 and GFP or expressed one of the two (Figure 3I-J). The difficulty with distinguishing temporal order in this assay suggests that both naive *Oct4-*GFP and DPPA4 are independently activated in a stochastic manner at a similar time.

Collectively, our results showed that naive-specific *Tet1-*mCherry reporter expression marks a distinct intermediate stage separating early and late phases of pluripotency transitions. The early phase is marked by the first detectable expression of endogenous *Nanog*, *Tet1* and *Oct4* in close sequence, and the late phase by activation of both *Dppa4* and the *Oct4* naive-specific distal enhancer (Figure 3K). This order of events is conserved during reprogramming in different media conditions and strain backgrounds tested in this study, using either male or female transgenic MEFs.

### Transcriptome changes in dual reporter-sorted reprogramming intermediates

Based on the FACS time-course profiles (Figure 3C-D), we selected the KSR+AA media condition to sort reprogramming intermediates from each dual reporter combination at the following time points: day 8, day 10 and day 12 for OKSM; *Tet1*^tm1.1Koh^; *Pou5f1*^tm2Jae/J^ (“totalx2”) and day 10, day 12 and day 14 for OKSM; Tg(*Tet1-*mCherry)B; TgOG2 (“naivex2”). At each time point, mCherry+/GFP− (mCh+GFP−) single-fluorescent and mCherry+/GFP+ (mCh+GFP+) double-fluorescent clusters were isolated by cell sorting, except in day 8 “totalx2” and day 10 “naivex2” samples in which mCh+GFP+ cells were too few to enrich (Figure 4A). A total of 10 sorted intermediate fractions, together with the parental MEFs and terminal iPSC lines clonally expanded for two early passages in serum and subsequently for another 4-5 passages in KSR, were analyzed using RNA-seq (n=3). Although the sequential order of reporter activation appeared independent of gender, that does not preclude that global transcriptomic and DNA demethylation kinetics may differ between male and female iPSCs during reprogramming ^41^. To avoid the gender confounder, we analyzed male lines only in these subsequent genomic analyses.

**Figure 4.**
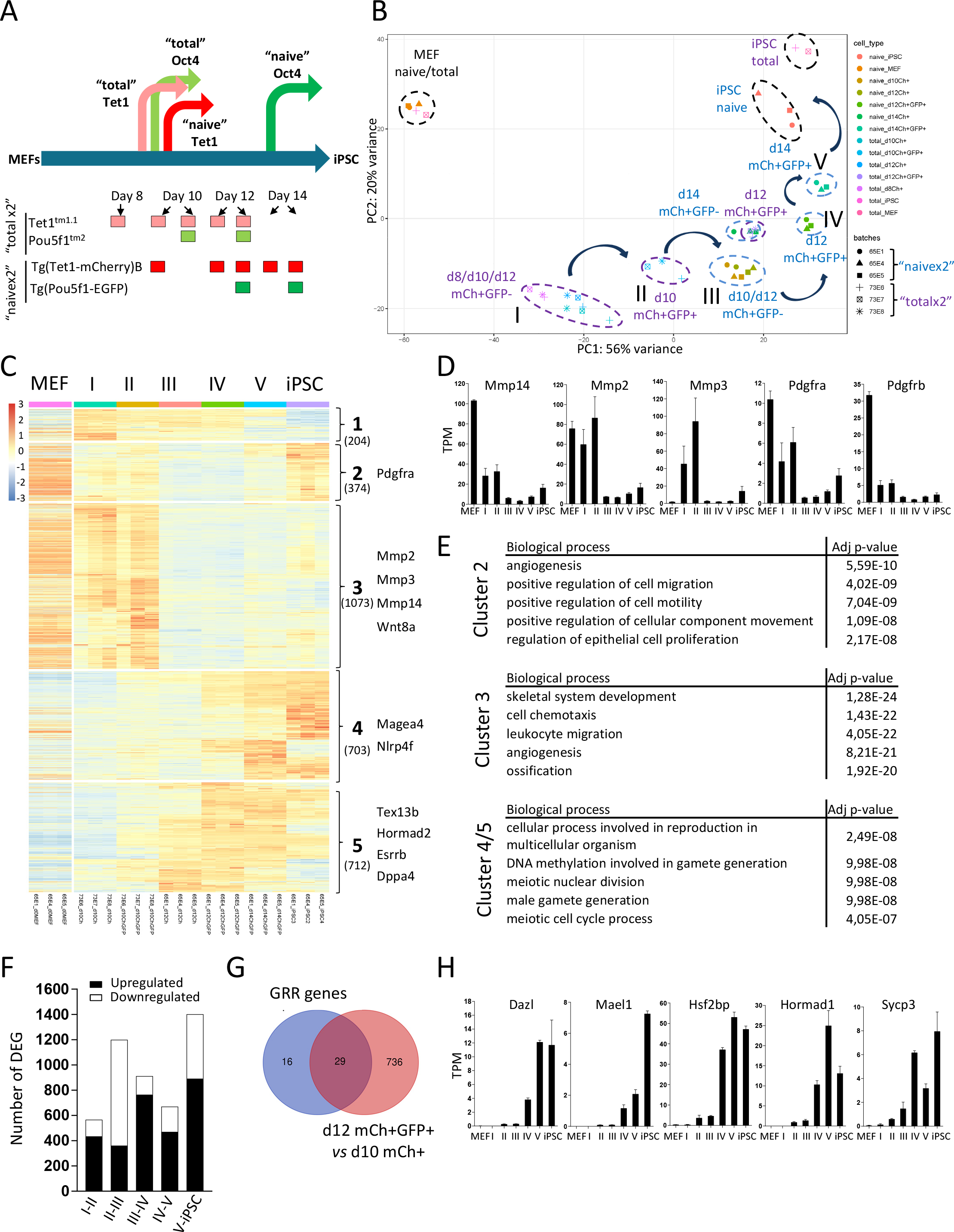
Transcriptome changes in dual reporter-sorted reprogramming intermediates. (A) Schematic of sorting strategy for Tg(*Tet1-*mCherry)B; Tg(*Pou5f1*-EGFP)Mnn (“naivex2”) and *Tet1*^tm1.1Koh^; *Pou5f1*^tm2Jae/J^ (“totalx2”) reporter lines. (B) Combined gene expression PCA plot of sorted populations of “totalx2” and “naivex2”, each in biological triplicates. MEFs and iPSCs are from the “totalx2” (n=2) and “naivex2” (n=3) lines. (C) Heat map of differentially expressed genes (DEGs) in pairwise comparisons between consecutive stages (Stage 1 to iPSCs) (adjusted *P* value < 0.05 and |log_2_ fold-change (FC)| > 2). MEFs are added in the left panel but were not included in the differential analysis. The number of genes in each cluster is shown in parenthesis. Representative genes are listed on the right. (D) Transcripts per kilobase million (TPM) of selected genes involved in migration and motility. Values are shown as mean ± SEM (n=3). (E) Top 5 Gene Ontology (GO) terms associated with gene clusters identified in Figure 4C. Clusters 4 and 5 are analyzed together. (F) Total numbers of DEGs upregulated or downregulated (adjusted *P* value < 0.05 and |log_2_ fold-change (FC)| > 1). (G) Overlap of genes upregulated from Stage III to IV with germline reprogramming-responsive (GRR) genes. See gene list in Figure S4G. (H) Same as 4D for selected GRR genes.

Principal component analysis (PCA) clearly separated the 10 intermediate fractions into 5 distinct groups, termed Stages I-V. These reflect a progression of transcriptional states along a trajectory towards the fate of clonally established iPSC lines (Figure 4B). Stage I comprised single-fluorescent mCh+GFP− fractions at all three time points of “totalx2” libraries. Next to these, the day 10 mCh+GFP+ triplicates separated as Stage II in the direction towards the iPSC clusters, supporting our live imaging assessment that the initial phase of *Tet1* gene activation precedes that of *Oct4*. By day 12, “totalx2” mCh+GFP+ samples coincided with “naivex2” mCh+GFP− samples at Stage III, which reflected initial activation of the *Tet1* naive-specific distal promoter, again consistent with a time-course trajectory in which activation of the *Tet1* distal promoter followed that of the *Oct4* promoter-proximal enhancer. This connection between “naivex2” and “totalx2” samples at Stage III also confirmed that the time-course trajectory we proposed is independent of strain differences between the two library preparations. Subsequent “naivex2” mCh+GFP+ samples collected at days 12 and 14, marking the closely related Stages IV and V respectively, were progressively closer to terminal iPSC lines, in line with *Oct4* distal enhancer activation being a late event in the maturation of iPSCs (Figure 4B). Notably, the samples clustered more closely by dual fluorescence patterns than by day of collection. Thus, the dual reporter system is a stringent way to isolate reprogramming intermediates from a cell population that is highly heterogeneous, i.e. they do not initiate reprogramming at the same time even if every cell started with the same stoichiometry of reprogramming factors.

To compare these stages with earlier ones previously studied, we compared our mRNA-seq datasets with a recent total RNA-seq by Knaupp et al. that examined reprogramming intermediates sorted based on cell surface THY1-/SSEA1+/C-KIT+ fractions from day 3 to day 12 ^11^. On the same PCA plot, our late iPSC intermediates clustered relatively closer to iPSCs compared to the Knaupp’s earlier intermediate samples; conversely, Knaupp’s samples were closer to MEFs (Figure S4A). This comparison confirms that our sorted late-intermediates represent novel stages reflecting a continuum of pluripotency remodeling events during the maturation phase of iPSC reprogramming.

We selected samples to represent each stage (I: “totalx2”_day 10_mCh+GFP-, II: “totalx2”_day 10_mCh+GFP+, III: “naivex2”_day 12_mCh+GFP-, IV: “naivex2”_day 12 _mCh+GFP+, V: “naivex2”_day 14 _mCh+GFP+, “naivex2” iPSC) for further analysis. We first performed pairwise comparisons between consecutive stages (Stage II versus I, Stage III versus II,…, iPSC versus Stage V) to compile all genes showing differential expression in at least one pairwise comparison. Using stringent filtering criteria (adjusted *P* value < 0.05 and |log_2_ fold-change (FC)| > 2), we identified a set of 3066 differentially expressed genes (DEGs) for hierarchical clustering (Figure 4C). The heat map revealed global waves of transcriptomic changes. These were most prominent between Stages II and III, corresponding with transition to early activation of the *Tet1* “naive” distal promoter, when a majority of genes dramatically lost expression. This transition was followed by predominantly gene activation at other loci during the subsequent Stages III to V, until the final transition to established iPSCs. Normalized read counts showed progressive induction of pluripotency genes beginning at Stage II (Figure S4B). Based on the stage at which iPSC levels of expression were attained, they can be classified into early (e.g. *Nanog*, *Prmd14* by Stage II), intermediate (e.g. *Esrrb* by Stage III and *Rex1*, *Tbx3* by Stage IV) and late (e.g. *Dppa4* by Stage V) factors. In contrast, primed-state markers such as *Wnt8a* were silent by Stage III (Figure S4B).

Using a hierarchical threshold of 5, we defined the gene clusters exhibiting distinct patterns of stage-dependent transitions (Figure 4C). Cluster 1 described genes that that transiently gained expression during the intermediate stages and were downregulated late in Stage V intermediates and iPSC clonal lines; this small set was not significantly enriched for any gene ontology (GO) terms. Cluster 2 comprised genes downregulated from Stage II to III but later restored in expression in iPSCs; these were significantly enriched in GO terms related to cell migration and motility; an example is *Pdgfra* (Figure 4C-E). Cluster 3 contained a major group of genes that were expressed in MEFs and until Stages I and II but downregulated upon activation of naive markers, although some were modestly up-regulated again in iPSCs; these genes were highly specific for processes related to development, structure organization and cell migration, including the matrix metalloproteinases *Mmp2*, *Mmp3*, *Mmp14* (Figure 4D-E). In contrast, Clusters 4 and 5 collectively comprised genes activated sequentially in association with naive-state activation of *Oct4* and *Tet1* distal elements respectively, including a subset with transient expression in the intermediate stages but not in iPSC lines. Genes in Clusters 4 and 5 were mostly involved in meiosis and gametogenesis (Figures 4C and 4E).

Stage III appears to be a pivotal branch point in the pluripotency trajectory and may be represented by either “naivex2” day 12 mCh+GFP− or “totalx2” day 12 mCh+GFP+ samples. To discern subtle differences between the two samples, we added the expression profiles of “totalx2” day 12 mCh+GFP+ samples in the heat map based on the DEG clusters defined in Figure 4C. On this composite heatmap (Figure S4C), “totalx2” day 12 mCh+GFP+ appeared as an intermediary state between Stages II and III, distinguished from “totalx2” day10 mCh+GFP+ mainly by lower expression of DEG clusters 2 and 3. That “totalx2” day 12 mCh+GFP+ clustered closer to Stage III than to Stage II on the PCA is consistent with our proposed trajectory, where these are cells that had progressed for another 2 days after first co-activation of total mCh+GFP+ and likely were beginning to activate the naive *Tet1* distal promoter, but could also be more heterogeneous. For further analysis, we chose to examine the route signaled by early activation of each of the four reporters in sequential order (Stage I to IV), which more likely marks the most efficient pathway.

To examine a more comprehensive set of DEGs, we performed pairwise differential expression analysis between consecutive stages using less stringent criteria (adjusted *P* value < 0.05, |log_2_FC| > 1). Assessment of these expanded set of DEGs revealed similar GO terms enriched as those shown in Figure 4E. Briefly, terms associated with specific stage transitions include pattern specification and cell fate commitment from Stage I to II, cell migration and motility from Stage II to III, meiotic and gamete specification from Stage III to IV and again cell migration and motility in Stage IV to V (Figure S4D). The transition from Stage II to III was unique by showing downregulation of two-thirds of DEGs, in contrast with the majority of up-regulated DEGs in the other intermediate stages (Figure 4F). Thus, the acquisition of naive pluripotency signatures during the late intermediate stages to mature iPSCs may involve a temporary stalling of cell proliferation and movement, which is restored once iPSC lines are clonally expanded.

To determine whether the same set of up-regulated DEGs accumulated over time, we compared all “naive” stages (IV, V and iPSCs) with the earliest stage at which the “naive” *Tet1-* mCherry reporter was active (“naivex2” day 10 mCh+GFP−). Indeed, we observed that 74% of DEGs (680 of 923) arising from the transition from single mCh+GFP− to dual mCh+GFP+ by day 12 remained differentially expressed by day 14, after which 65% (1094 of 1676) persisted as DEGs until the iPSC stage, reflecting stable gain or loss of expression during iPSC maturation (Figure S4E-F). We also observed that 29 genes differentially expressed between day 10 Ch+GFP− and day 12 Ch+GFP+ were among 45 recently identified germline reprogramming-responsive genes (GRRs) ^42^. The latter are marked by high-CpG promoters that became demethylated during germline epigenetic reprogramming concurrently with transcriptional activation in PGCs (Figures 4G-H and S4G).

### Methylome changes in reprogramming intermediates

Epigenetic reprogramming in the germ-line is known to involve a sequential two-stage process in which an early passive global wave of DNA demethylation is followed by locus-specific active demethylation ^43,44^. Therefore, we asked whether phases of stage-specific DNA demethylation may also be identified in our sorted intermediates. First, we analyzed by Q-PCR gene expression changes of TET DNA oxygenases and DNA methyltransferases (DNMT) during the different stages. Both *Tet1* and *Tet2* were expressed at intermediate levels at early Stage I, in line with early activation of the *Tet1* proximal promoter, and at full levels upon activation of the naive *Tet1* distal promoter (Stage III) (Figure S5A). This step-wise increase in *Tet1* and *Tet2* expression was associated with a reciprocal step-wise loss of *Tet3* expression, such that *Tet3* was silenced to basal levels once the naive *Tet1* distal promoter was fully active. Genes encoding the maintenance methylase *Dnmt1* and the associated protein ubiquitin-like with PhD and ring finger domains 1 *Uhrf1* were constitutively expressed throughout these stages. *Stella* (also known as *Dppa3*, *Pgc7*), which protects the genome from TET3 oxidation in the mouse zygote ^45^, was transiently upregulated from Stage II to V (Figure S5B). Expression of the *de novo* DNA methyltransferases varied more; *Dnmt3a* was constitutively expressed, undergoing a transient and modest downregulation during Stages III-IV before elevated expression in iPSCs while *Dnmt3b* was up-regulated progressively during the intermediate stages from low starting levels in MEFs (Figure S5C). These results suggested active and dynamic remodeling coupled with protection of the DNA methylome in the final stages of iPSC generation.

Since both *Oct4* DE and PE are known to undergo DNA demethylation during iPSC formation^1^, we examined methylation changes at the two loci in the sorted intermediates. Cell populations were sorted at day 10 from “totalx2” and at day 14 from “naivex2” dual reporter systems. Compared to the highly methylated status of CpGs at the *Oct4* promoter and DE in MEFs and in day 14 double-negative cells that failed to reprogram, we noted that the earliest evidence of *Oct4* promoter demethylation occurred in day 10 “totalx2” mCherry+/GFP+ populations, in line with detection of the total *Oct4-*GFP reporter (Figure 5A). Promoter demethylation was maintained in the “naivex2” day 14 reprogramming intermediates (Figure 5B). On the other hand, methylation erasure of the *Oct4* DE was evident later only in day 14 “naivex2” mCherry+/GFP+ populations (Figure 5A-B), in line with naive-state reporter activation. Of note, a distal site near *Tet1* exon 1b already showed partial loss of methylation in sorted cells from day 10 “totalx2” and became fully demethylated in day 14 “naivex2” fractions in concordance with mCherry expression. Thus, the DNA methylation status at each site strictly follows reporter activation kinetics of the target gene.

**Figure 5.**
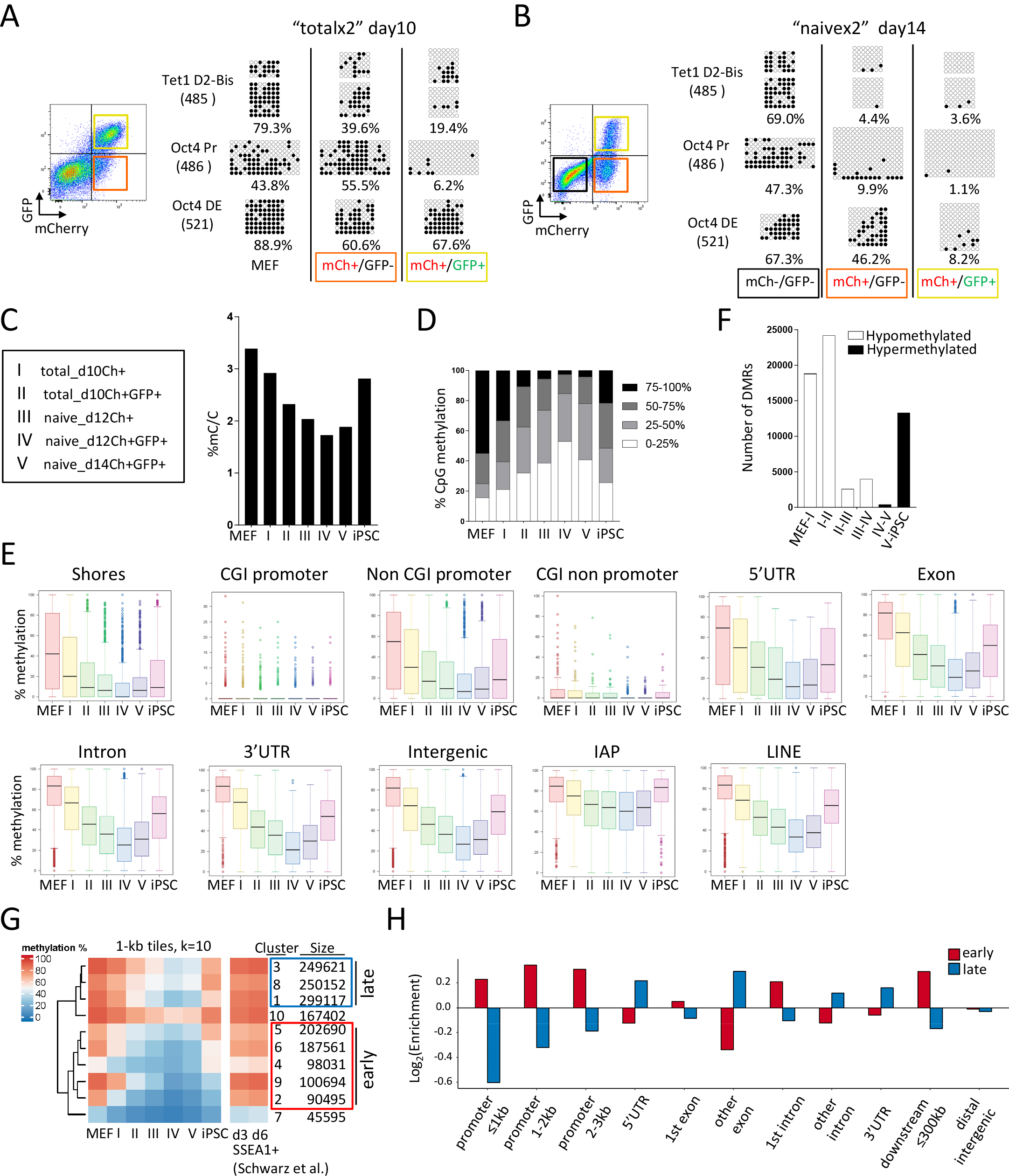
Methylome changes in reprogramming intermediates. (A-B) Bisulfite sequencing of *Tet1* distal promoter (D2-Bis); *Oct4* proximal promoter (Pr) and *Oct4* distal enhancer (DE) loci. Left panels: flow cytometry plots highlighting the sorted populations from Tg(*Tet1-*mCherry)B;Tg(*Pou5f1*-EGFP)Mnn line (A) or Tet1tm^1.1Koh^;*Pou5f1*^tm2Jae/J^ line (B). Colored squares indicate mCh+GFP− (red), mCh+GFP+ (yellow), mCh-GFP- (black) populations (only in B). Right panels: CpG methylation profiles of *Oct4* loci (bp sequence length in parenthesis) compared with MEFs (A) or mCh-GFP- (B). Closed circles denote 5mC (or 5hmC), open circles unmethylated C; X indicates a missing sequence or mismatch. Values indicate average methylation levels (%) of clonal sequences shown. Results are representative of 2 independent experiments. (C-H) Oxidative bisulfite sequencing (true 5mC) at genome-scale of MEFs, reprogramming intermediates and iPSCs. (C) Percentage of methylation calculated as (CG+CHH+CHG)/total C’s (whereby H can be either A,T or C). Legend on the right explains samples representing each stage. (D) Distribution of CpGs classified by methylation levels (25% bins). See Figure S5D for correlation and full histogram plots. (E) Box-whisker plots of CpG methylation levels (base coverage ≥10) across genomic features. (F) Differentially methylated regions (DMRs) classified as hypermethylated or hypomethylated in reprogramming intermediates. (G) Heatmap of clustered 1-kb tiling regions. Clustering was based on k-means clustering with k = 10. Clusters classified as early and late waves of demethylation are marked by red and blue boxes respectively. (H) Genomic features enriched in tiles associated with early and late waves as compared to all tiles from Figure 5G.

To define globally the methylome changes at each intermediate stage (I to V) compared to MEFs and iPSCs, we performed whole genome bisulfite sequencing (WGBS) following oxidative bisulfite conversion ^46^. By this approach, we detected true 5mC in distinction from 5hmC at base-resolution, of which >80% are positioned within CpG dinucleotides at all intermediate stages. The global content of mapped methylated cytosines decreased progressively from 3.4% in MEFs to 1.7-1.9% by Stages IV and V, before regaining in clonal iPSCs (Figure 5C). At all CpGs covered by 10 or more reads (comprising > 90% CpGs sequenced), we calculated the distribution of CpG methylation levels in each sample. As previously shown, MEFs showed bimodal distribution peaks contributed by lowly (0-25%) and highly methylated (75-100%) CpGs ^47^ (Figure 5D and S5D). In iPSCs, the fraction of highly methylated CpGs was reduced, whereas the lowly and intermediately methylated fractions increased. Intermediates stages showed a progressive loss of highly methylated CpGs, reaching the most globally hypomethylated state by Stage IV, when both *Tet1* and *Oct4* distal elements are active. Further progression and clonal expansion of iPSCs were associated with *de novo* DNA methylation that re-established a subset of highly methylated sites, but the global methylation status of iPSCs remained hypomethylated relative to parental MEFs (Figure 5D and S5D).

We then analyzed all CpGs with a base coverage exceeding 10-fold across all samples. In consecutive Stage I to V intermediates starting from MEFs and terminating as iPSCs, we observed a progressive wave of demethylation that appeared global, sweeping occurring across CpG island (CGI) shores, non-CGI promoters, 5’UTR, exons, introns, 3’UTR and intergenic regions (Figure 5E). As previously shown in PGCs ^48^, also repetitive sequence classes such as long interspersed nucleotide elements (LINE) showed demethylation reprogramming, but the class of parasitic retrotransposon intracisternal A particle elements (IAPs) were largely resistant (Figure 5E). As a separate class, the majority of CGIs were unmethylated across all cell stages within or outside gene promoters, although a minor subset of highly methylated CGIs in MEFs showed methylation loss during progression through the reprogramming stages. At known germ-line imprinting control regions (ICRs) ^49^, where parental allele-specific differential methylation regions (DMRs) are preserved from the germ-line during development, we calculated average methylation levels of CpGs per locus (base coverage ≥ 5). Since our datasets lack allele-specific methylation information, we assessed whether intermediate (~25-75%) methylation levels persisted at these loci. In general, intermediate levels of methylation were variably eroded at several loci across stages, most notably at the paternal imprint locus *Gpr1/Zdbf2* and the maternal loci *Mest/Peg1* and *Snurf/Snrpn* at Stages IV and V (Figure S5E).

We further defined DMRs in pairwise comparison between consecutive stages. The bulk of global CpG demethylation progressed early during Stages I and II (Figure 5F). From Stage III to IV, further loss of CpG methylation was more gradual despite full expression levels of *Tet1* and *Tet2* in the naive state (Figures 5F and S5A). Overall, the kinetics of DNA methylation reprogramming suggests an early major global wave of demethylation upon entry into pluripotency, which coincides with early gene activation of endogenous *Tet1* and *Oct4*, followed by more gradual demethylation when naive pluripotency features are acquired.

To better define these sequential waves of global demethylation, we calculated CpG methylation levels across non-overlapping 1-kb tiles genome-wide and collated a set of 1.7 × 10^6^ tiles that fulfilled our coverage threshold (see Methods) in all samples analyzed. Collectively, these covered 62.1% of the genome and 91.1% of all promoters. Unsupervised k-means clustering of all tiles indeed suggests two general waves of global DNA demethylation: an early wave (Clusters 5+6+4+9+2) that was completed by Stage II and a later wave (Clusters 3+8+1) that reached the hypomethylated state after Stage III (Figure 5G). Collectively, both early and late cluster waves impacted 87.4 % of the covered genome. These losses of 5mC methylation were not apparent in day 3 and day 6 SSEA1-sorted intermediates examined in a previous study ^50^ which showed an early wave of DNA demethylation occurring at a subset of ESC enhancers (Figure 5G).

Genomic annotation of both cluster groups representing early and late demethylating waves revealed an association with distinct features. The early wave occurred at regions enriched in promoters, 1^st^ introns, and downstream (≤300 kb) regions and depleted within gene body regions, relative to the late wave where 5’UTR, other exons, other introns and 3’UTR were enriched (Figures 5H and S5F). Reclassification of these regions by ESC functional annotation showed an enrichment of the early wave at active and poised promoters and enhancers and depletion at transcriptional (txn) elongation sites, where instead the late wave is reciprocally enriched (Figure S5G-H). These observations validate the kinetics of global DNA demethylation during iPSC reprogramming as a two-stage process that targets predominantly promoters and upstream elements early and then the remainder of gene bodies later.

To understand how these DNA demethylation waves relate to transcriptomic changes at target genes, we calculated CpG methylation levels across promoter regions (−2kb to +0.5 kb of TSS) of genes depicted in the DEG clusters of Figure 4C. In all five DEG clusters, non-CGI promoter methylation were synchronously erased by Stage II, independently of the differential kinetics of gene expression changes in each DEG cluster (Figure S5I). These observations clearly indicate an uncoupling of DNA methylation and gene expression changes at these late intermediate stages of reprogramming.

### Reprogramming in the absence of TET1

The temporal sequence by which naive-state *Tet1* gene activation preceded that of *Oct4* suggests a deterministic function of TET1 in full acquisition of naive pluripotency in iPSCs. However, previous studies using MEFs in which *Tet1* was disrupted by gene targeting nonetheless generated iPSC colonies that could even contribute to blastocyst chimera ^10^; only knockout of all three TET genes disrupted reprogramming at the early stage of mesenchymal-to-epithelial transition ^51^. In these previous studies, the *Tet1* gene was inactivated respectively by deleting exon 5 or exons 11-13 to disrupt expression of the catalytic domain. Because these earlier genetic knockouts (KOs) of *Tet1* may not have disrupted non-catalytic functions provided by the N-terminal domain of TET1, we assessed reprogramming using *Tet1* KO MEFs generated from our B6-*Tet1*^tm1Koh^ strain, which disrupts *Tet1* expression using a knock-in (KI) cassette after the ATG start codon ^27^. We crossed the *Tet1*^tm1Koh^ allele to the TgOG2 strain to assess whether naive-state activation of *Oct4* DE as reflected by the GFP reporter could be affected by complete loss of TET1.

Wild type (wt) and KO MEFs containing the naive *Oct4-*GFP reporter were harvested from littermate timed-pregnant embryos for reprogramming in parallel using lentiviral OKSM transduction, comparing cells of the same gender. Cells were collected every other day for FACS and stained with SSEA1 antibody to identify the population that would enter the pluripotent pathway towards activation of *Oct4-*GFP ^8^. By day 7, ~60% of both wt and *Tet1* KO cells started to express SSEA1. These SSEA1+ populations then shifted over time to gain also *Oct4-*GFP, without any evidence of delay in the KO compared to wt in both male and female pairs (Figure S6A-C) in either serum or KSR media conditions in the absence or presence of ascorbic acid. Final counts of alkaline phosphatase-positive colonies were also not affected by absence of TET1 (Figure S6D). We verified absence of *Tet1* coding transcripts in KO clonal iPSCs and in cells harvested during the time-course with two set of Q-PCR primers detecting exons 3-4 and exons 12-13; we found no evidence that alternative transcripts were produced from downstream TSSs in any detectable amounts (Figure S6E-F). RNA-seq IGV tracks also showed absence of reads across the entire *Tet1* coding sequence in *Tet1* KO iPSCs (Figure S6G).

As *Tet2* is also upregulated during reprogramming in parallel with *Tet1* (Figure S5A), we depleted *Tet2* expression in *Tet1* KO MEFs using two independent lentiviral shRNA knock-down (KD) constructs during lentiviral OKSM reprogramming. In these *Tet1* KO; *Tet2* KD reprogramming cells, we observed activation of “naive” *Oct4-*GFP nonetheless (Figure S6H-J). Demethylation of the *Oct4* DE has previously been observed in reprogramming from *Tet1; Tet2* double KO MEFs ^10^, suggesting that methylation erasure at this locus may be passive.

To determine if *Tet1* KO iPSCs may nevertheless display epigenetic defects, we profiled the transcriptome and methylome of clonal lines of *Tet1* KO iPSCs in comparison with wt cells reprogrammed from littermate controls. PCA analysis of RNA-seq data from clonal wt and KO iPSC clonal lines showed both wt and KO to cluster close together where terminal iPSCs were expected, away from all other reprogramming intermediates (Figure 6A). Thus, reprogramming can progress to gain terminal iPSC status in the absence of TET1, albeit possibly by an alternative route.

**Figure 6.**
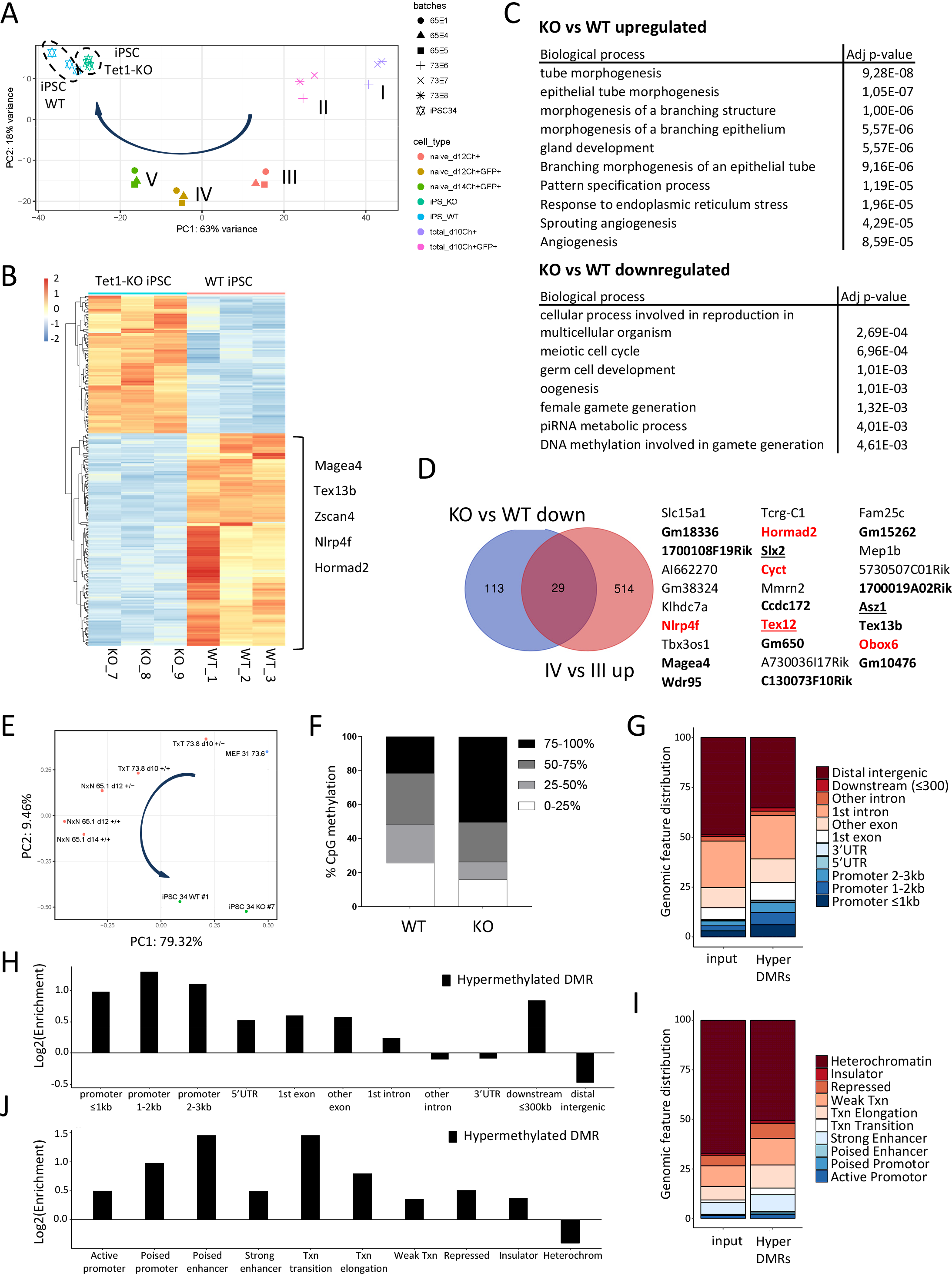
Reprogramming in the absence of *Tet1*. (A) Gene expression PCA of the 5 intermediates from Figure 4C compared to wt and *Tet1-*KO iPSCs (circled with a dotted line), in biological triplicates. (B) Heat map clustering of DE genes in wt and *Tet1-*KO iPSCs. (C) GO terms associated with genes up-regulated (top 10) and down-regulated (all) in *Tet1-*KO iPSCs compared to wt. (D) Overlap of down-regulated genes in *Tet1-*KO compared to wt iPSCs with genes upregulated during Stage III to IV transition. The list of common genes is shown on the right. Genes in red are analyzed in Figure S7. Genes showing germ cell-restricted expression are in bold; GRR genes are underlined. (E) PCA of 5 intermediates including WT#1 and KO#7 iPSC lines based on methylation level of 1-kb tiling regions from Figure 5G. Arrows indicate the direction from MEFs to iPSCs. See Figure S6L for PCA based on single CpG methylation differences, including replicate wt and KO samples. (F) Distribution of CpGs classified by methylation levels in WT#1 and KO#7 iPSC. (G-H) Distribution of genomic features (G) and their enrichment in hypermethylated DMRs in *Tet1-*KO compared to wt (H). (I-J) Same as G-H for ESC specific functional features.

To gain insight into phenotypic differences between wt and KO iPSCs, we examined differential gene expression between the two groups. Heat map analysis of DEGs revealed a larger proportion of genes downregulated in the KO compared to the wt, consistent with gene silencing caused by loss of promoter demethylation in the KO (Figure 6B). GO analysis of upregulated genes in KOs described a broad variety of morphogenic processes. In contrast, downregulated genes enriched for pathways specifically related to gametogenesis and meiosis and include some genes expressed exclusively in germ cells such as *Magea4*, *Tex13b* and *Hormad2* (Figure 6B-C). Furthermore, the entire *Zscan4* family gene cluster, required for telomere homeostasis and karyotypic stability of ESCs in self-renewing cultures ^52^, was down-regulated in KO iPSCs (Figure 6B). The latter is reminiscent of our recent observations in *Tet1* KO EpiLCs, where similar downregulation of *Zscan4* family genes resulted in accumulation of cells with shortened telomeres ^27^.

The DEGs up-regulated in KO iPSCs relative to wt showed little overlap with those down-regulated during Stage II to III transition, suggesting that loss of TET1 did not affect the global gene repression observed at Stage III, possibly due to functional redundancy with TET2 which was still expressed in TET1-deficient iPSCs. However, both DEGs downregulated in KO iPSCs and those up-regulated during transition from Stage III to IV were similarly enriched for GO terms in meiosis and gamete generation, suggesting that TET1 may be directly responsible for activating these genes. Indeed, we identified 29 genes common in both sets of DEGs in the Venn diagram overlap (Figure 6D). Many of these have no known functions, but 18 are documented to show germ cell-restricted expression in GEO expression databases, including *Magea4*, *Tex13b, Tex12*, *Asz1* and *Slx2* (Figure S6K). The last three are also GRR genes (Figure 6D).

To compare the methylome of wt and KO iPSCs, two replicate sample pairs generated from different strain crossings (estimated 62.5% B6; 12.5% CBA; 25% 129S6 or 75% B6;25% CBA) were examined by oxidative WGBS. Similar to the RNA-seq analysis, the methylome PCA showed KO iPSC lines acquiring a methylome status closer to wt iPSCs than to the intermediates (Figure 6E and S6L). However, high coverage analysis of CpG methylation levels in a pair of wt and KO iPSC lines showed a striking re-distribution of methylation frequencies towards an increased proportion of fully methylated CpGs in KO compared to wt (Figure 6F and S6M). Furthermore, we identified 5,495 DMRs with more than 25% methylation level differences between wt and *Tet1* KO samples, of which 5,492 were hypermethylated and only 3 hypomethylated in the KO relative to wt. Hypermethylated DMRs were enriched within promoters but not at distal intergenic regions, and also within active regulatory elements but depleted at heterochromatin regions, suggesting that there could be a functional impact on gene expression (Figure 6G-J). These results are consistent with a critical role of TET1 in active DNA demethylation during reprogramming, loss of which generates iPSCs with grossly aberrant DNA methylation patterns.

We asked whether there are specific genes that dependent on TET1 for both induction of promoter DNA demethylation and gene expression during reprogramming. Of the 29 DEGs that were upregulated late during Stage III-IV reprogramming (Figure 6D), we examined five (*Tet12*, *Nlrp4f*, *Obox6*, *Hormad2* and *Cyct*) in further detail (Figure S7A-C). These five genes exhibited erasure of 5mC at promoter regions by Stage IV in *Tet1* expressing cells, but regained 5mC in KO iPSCs (Figure S7B). Q-PCR expression analysis of these genes in SSEA1+-sorted cells during reprogramming verified that they were all down-regulated in KO cells by day 13, with sustained loss of expression in KO iPSCs (Figure S7C). Thus, a subset of genes strictly depends on TET1 for promoter demethylation and gene activation late in reprogramming.

Overlapping the full set of TET1-dependent DMRs in iPSCs with both early and late demethylating regions identified in the stage intermediates suggests an active role of TET1 in both waves but with distinct biological functions. DMRs that appeared in the early wave were enriched in GO biological process related to differentiation, biosynthesis and signaling (Figure 7A). In contrast, the top GO term enriched by DMRs in the late wave was DNA repair (Figure 7A). Since KO iPSCs also showed loss of *Zscan4* expression, we examined telomere integrity in wt and KO iPSCs using a flow cytometry-based fluorescent *in situ* hydridization (flow-FISH) assay to quantify telomere lengths. Based on our previous study that showed that telomere defects are apparent in *Tet1* KO cells upon primed epiblast differentiation but not in naive ESC cultures ^27^, we converted our wt and KO iPSC lines to EpiLCs. Under this lineage-primed state, we observed an increased fraction of sub-diploid cells in KO lines compared to wt (Figure 7B). In cells gated in the G1 cell cycle phase, the telomere fluorescent probe detected increased fractions of cells with shortened telomeres in KO compared to wt (Figure 7C). These observations are suggestive of impaired telomere homeostasis in KO cells and consistent with regulation of chromosomal stability by *Tet1-*mediated DNA demethylation during reprogramming.

**Figure 7.**
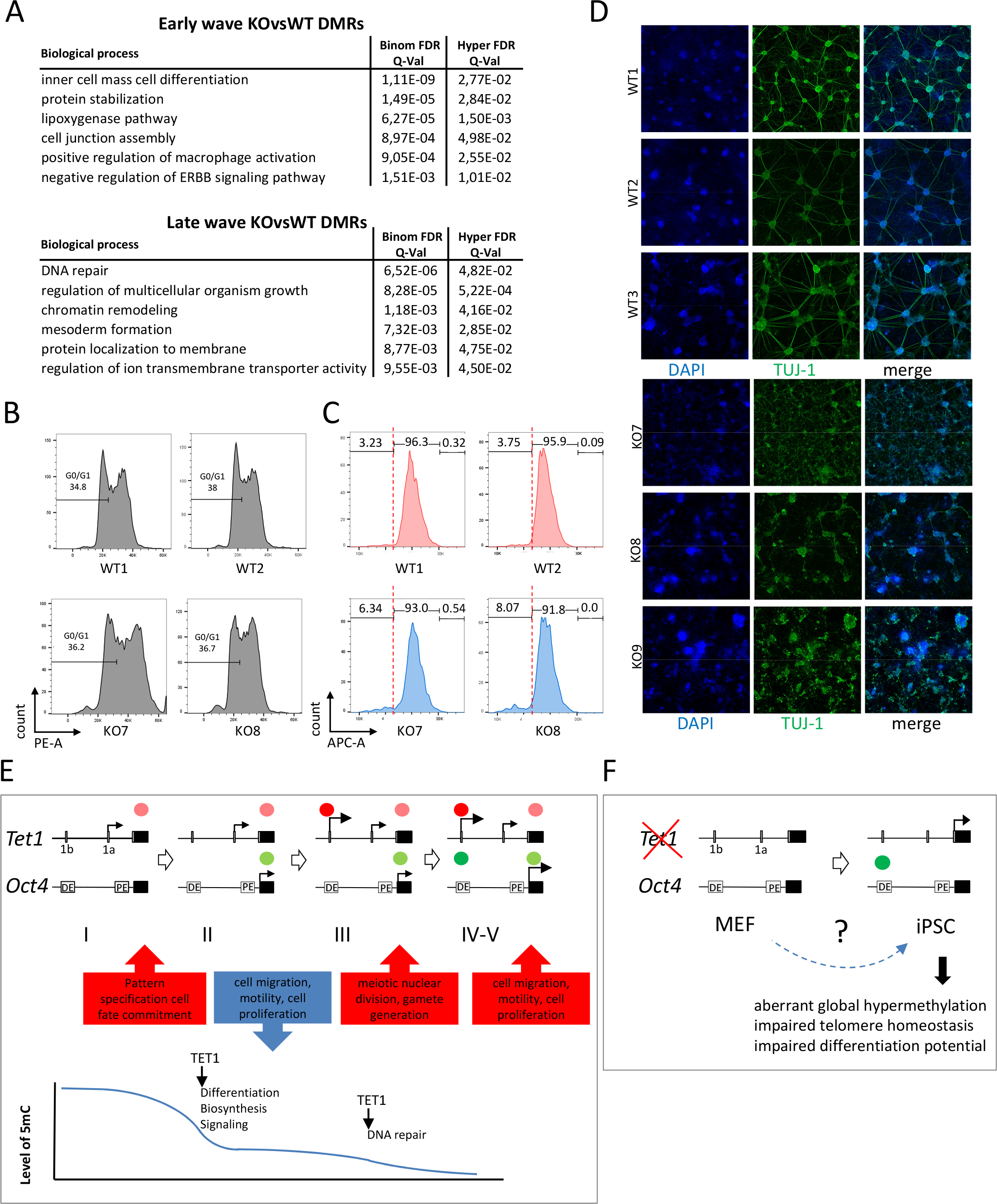
Differences between wt and *Tet1-*KO iPSCs. (A) Top 6 GO terms selected based on both binomial and hypergeometric FDR Q-values < 0.05 and number of observations > 5 related to genes nearest to *Tet1-*dependent DMRs overlapping with early and late demethylation clusters identified in Figure 5G. (B-C) Flow-FISH analysis of wt and *Tet1-*KO EpiLC lines (biological duplicates) showing cell cycle analysis by propidium iodide staining (B) and telomere length analysis using a telomere probe (C). Cells in C are gated in the G1 phase as shown in B; percentage values are shown for fractions gated by telomere length. Left of dashed lines mark cell fractions with sub-telomere length. (D) Immunofluorescence staining for TUJ-1 of wt and *Tet1-*KO iPSCs (n=3 biological replicates) during neuronal differentiation. (E) Schematic overview to illustrate how the dynamics of *Tet1* and *Oct4* gene activation identify temporally distinct late stages in reprogramming. (F) Schematic representation of an unknown (?) alternative trajectory taken by TET1-deficient cells, generating iPSCs of compromised quality.

Finally, we asked whether these epigenetic defects in KO iPSCs affected their differentiation potential. Using a terminal neuronal differentiation assay, we observed efficient generation of TUJ-I positive neurons using our wt iPSCs. In stark contrast, KO iPSCs differentiated poorly into TUJ-I positive cells, forming disorganized cell clusters and synaptic networks (Figure 7D). While we cannot rule out a direct role of TET1 in neuronal differentiation, these results suggest that epigenetic defects in KO iPSCs are associated with compromised differentiation potential.

## Discussion

In this study, we used dual reporters reflecting both naive-state specific and total expression of *Tet1* and *Oct4* to define a sequential order of reprogramming in the final phases of iPSC generation. Soon after detection of NANOG expression, which marks the gateway into pluripotency ^10,13^, reprogramming follows a one-way trajectory involving sequential activation of *Tet1* and *Oct4* proximal regulatory elements (Stages I and II) and a subsequent activation of distal promoter/enhancer regions (Stages III and IV), in which naive-specific *Tet1* gene activity clearly precedes that of *Oct4*. Using these dual reporter patterns as stage markers, we captured successive waves of gene expression and methylation reprogramming leading to the establishment of iPSC lines (Figure 7D). Dysfunction in TET1-mediated DNA demethylation during reprogramming, despite having no effect on activation of endogenous *Oct4*, causes epigenetic abnormalities, impaired telomere homeostasis and loss of differentiation potential in iPSCs (Figure 7E). Importantly, our results suggest that proper *Tet1* expression during reprogramming is an intrinsic component of an epigenetic rewiring program essential for ultimately generating high quality and developmentally potent iPSCs.

In two previous studies using doxycycline-inducible secondary murine reprogramming systems to track the epigenetic pathways towards induced pluripotency, cells were collected as heterogeneous unsorted samples at defined time points, but omitting the time window between day 11 and 16 when iPSCs were starting to form ^14,53^. Another study using clonal analysis in late stages focused on sample collection after day 14 ^54^. Although time-points between studies conducted using different systems cannot be directly compared, the consensus knowledge from these studies provided limited temporal resolution in the late stages. Here, we sorted reprogramming intermediates between days 8 to 14 using our *Tet1/Oct4* dual reporter iPSC system in sufficient amounts for genomic analysis. This approach efficiently isolated cells synchronized at defined cell state transitions based on gene reporter patterns and therefore, minimized stochastic effects due to different reprogramming rates even in isogenic cells that undergo the same inductive period.

In comparison to other recently published studies that sorted earlier reprogramming intermediates until day 12 ^11,50^, our late-intermediate stages are clearly distinct in gene expression and methylation patterns, filling in a developmental continuum between days 10 to 14 prior to clonal expansion of iPSC lines. This reprogramming window is challenging to capture because successful reprogramming occurs typically in fewer than 0.05% of starting MEF numbers in conventional assays, such that formative iPSCs are difficult to isolate without clonal expansion. Although highly efficient reprogramming systems using C/EBPα-enhanced reprogramming of B cells and compound-accelerated reprogramming of fibroblasts into iPSCs have recently been described ^55^, the kinetics of these assays (within 4 days) are too rapid to depict progressive events after initial acquisition of pluripotency and likely captured only a subset of DNA methylation remodeling. Our study has captured major waves of DNA demethylation that covered more than half the genome.

In early mammalian embryogenesis, genome-wide erasure of DNA methylation occurs in the pre-implantation embryo and in PGCs of the early embryonic germ line ^43,49,56^. As part of a normal developmental program, such global reduction of DNA methylation appears disconnected from transcriptional regulation, suggesting alternative or compensatory mechanism of gene expression regulation to prevent aberrant transcriptional activation ^5^. In PGCs, a two-stage DNA demethylation program ensures proper gametogenesis ^57^. The first stage involves global loss of methylation in a passive manner during PGC expansion and migration (between E8.0 to E9.5), during which imprints, germline-specific and meiotic genes are actively protected. The second stage occurs upon entry of PGCs into the gonads (from E10.5), when those aforementioned genomic regions then undergo locus-specific demethylation requiring TET1 and TET2 ^44^. In contrast, protection of imprint-specific DMRs from global demethylation in the pre-implantation embryo is essential for proper embryonic development ^58^.

The genome-wide demethylation events captured in our experimental induction of pluripotency bear both remarkable similarities and differences with those occurring in physiological reprogramming. We observed a major wave of global DNA demethylation occurring early during Stages I and II. This progressed into a second late wave signaled by full *Tet1* gene activation (Stage III) to reach the most hypomethylated state (Stage IV) when *Oct4* DE is activated. The latter wave coincided with activation of late pluripotency and germline genes. Thus, two global waves of DNA demethylation can be temporally separated by an intermediary Stage III marked by activation of a naive-specific *Tet1* distal promoter, bearing a resemblance to germline epigenetic reprogramming. However, despite the stage-specific waves of expression changes in gene clusters that suggest a phenotypic transition occurring progressively at every stage, the global DNA demethylation is predominantly targeted at non-CGI promoter regions almost synchronously at Stage II across all gene clusters, suggesting an uncoupling of DNA demethylation and transcriptional regulatory mechanisms characteristic of early development. These results provoke a paradigm shift in understanding the maturation stage of iPSC generation, in that pluripotency genes are not immediately activated as a result of global DNA demethylation. Unlike PGC development, meiotic and germline-specific gene promoters did not appear to be protected from the early wave of global DNA demethylation. Rather than affecting distinct functional gene clusters, the two-stage process of genome-scale DNA demethylation in iPSC reprogramming occurs sequentially as an early wave that removes 5mC predominantly from promoter regions and a later wave that removes 5mC from gene bodies.

The unique dynamics of DNA demethylation and germline-specific gene expression during robust iPSC reprogramming predicts then that genomic imprinting may be subjected to unprotected erasure. As previously observed during reprogramming of mouse iPSCs ^59^, we observed variable 5mC loss in a subset of germline ICRs. At some ICRs (*Gpr1/Zdbf2* and *Grb10*), 5mC levels were increased to intermediate levels upon clonal iPSC expansion, suggesting that iPSCs may nonetheless retain some cellular machinery to restore imprint patterns. Stochastic erosion of imprint DMRs has been observed in a large-scale analysis of human iPSC lines, raising concerns that all iPSC lines should be screened for proper imprinting prior to any application in disease modeling and cellular therapy ^60^.

Are the global waves of DNA demethylation in reprogramming a result of active or passive processes? Our RNA-seq analysis did not detect any repression of genes encoding the maintenance DNA methylation machinery (*Dmnt1* and *Uhrf1*) or overall loss of *de novo* DNMTs (*Dnmt3a* and *Dnmt3b*) throughout the intermediate Stages I to V, although it remains to be tested whether nuclear exclusion of the proteins had occurred. Of note, *Tet3* is still expressed at Stage I at only slightly reduced levels relative to that in parental MEFs, and is only reduced to basal levels by Stage III when both *Tet1* and *Tet2* are highly expressed, such that at least one TET family gene is expressed at every stage of reprogramming. Thus, an active role of TET3 in mediating global DNA demethylation during Stages I and II is possible, in a similar way that maternal TET3 contributes to active demethylation of the early zygotic genome ^56,61^. After Stage III, the progressive up-regulation of a small subset of pluripotency and germline genes may involve loci-specific DNA demethylation by TET1 and TET2, analogous to the late wave of demethylation in PGCs ^44^. However, TET1 is already involved in promoter demethylation in the first global wave. Thus, successful generation of iPSCs may involve repurposing of pathways underlying epigenetic remodeling in both the mammalian pre-implantation embryo and PGCs, resulting in unique molecular dynamics that account for epigenetic differences between embryo-derived and *in vitro* experimentally generated iPSCs.

Clonal expansion of iPSCs is associated with re-gain in expression of cellular proliferative genes that were down-regulated earlier at Stage III (DEG cluster 2), suggesting that a transient slow-down in cellular proliferation and motility accompanies sequential activation of late naive pluripotency in the final phases of iPSC establishment. We observed the lowest methylation status in the preceding Stage IV (day 12) intermediates, after which *de novo* DNA methylation is partially re-established in iPSC lines which were clonally expanded. The partial restoration of DNA methylation in established iPSC lines may reflect a biological phenotype distinct from that of formative intermediates, or an acute adaptation to a culture media switch following colony picking. Thus, previous studies comparing clonally expanded iPSC lines with early intermediates and parental somatic cells may have under-estimated the extent of global DNA methylome reprogramming at late-intermediate stages masked by serum-culture induced re-methylation. Changes in transcriptome and methylome profiles may continue after clonal expansion of iPSC lines. In particular, long term passaging of iPSCs has been reported to result in erosion of random aberrant methylation and tissue-of-origin methylation memory ^62^. In view of these experimental variabilities in the propagation and maintenance of established iPSC lines, our analysis of late stage intermediates offers a more precise view of the reprogramming epigenetic landscape independent of post-iPSC assay culture factors. To exclude methylation differences caused by gender, we have examined DNA methylome reprogramming only in male lines. Gender-dependent differences in female lines will be of interest to study in future applications of our dual reporter systems.

Since up-regulation of *Tet1* and *Tet2* expression is one hallmark of iPSC formation, the precise functions of DNA dioxygenases in reprogramming have been studied previously by various groups, but with conflicting results. Earlier studies, using primarily shRNA knockdown approaches, observed reduction in iPSC colony counts upon *Tet1* and/or *Tet2* depletion, suggesting that both genes contribute to reprogramming efficiency ^63,64^. One study suggested that TET1 could even replace OCT4 in the reprogramming factor cocktail and is critical for active DNA demethylation of endogenous *Oct4* and other pluripotency gene loci ^65^. In contrast, subsequent studies using genetic knockouts showed that MEFs with single gene targeted deletion of *Tet1*, *Tet2*, or *Tet3*, or combined double TET deletions can nonetheless be reprogrammed to form iPSC colonies, although epigenetic defects in these mutant lines were not investigated further ^10,51,66^.

Our studies provide a clearer view of TET1 function in reprogramming. First, TET1 (and likely also TET2) is not necessary to demethylate endogenous *Oct4*. Second, *Tet1* expression at maximal pluripotency levels, signaled by activation of its naive-specific distal promoter at Stage III, is associated with global down-regulation of gene expression from Stage II, suggesting that TET1 may contribute to global gene repression independent of DNA methylation changes. The latter is in agreement with non-catalytic functions of TET1 in development ^27^. Finally, MEFs can be reprogrammed to iPSCs in the absence of TET1, but nonetheless harbor widespread DNA hypermethylation marks and gene expression differences from wt iPSCs, which can greatly compromise the phenotype of differentiated derivatives. Our observation of telomere loss in cellular fractions of KO, but not wt, iPSCs, at the lineage-priming stage of differentiation provocatively suggests that TET1 may regulate cellular senescence and be a critical factor in the rejuvenation process of iPSC reprogramming.

Thus, the canonical role of *Tet1* in active DNA demethylation is dispensable for iPSC generation, yet is critical for proper epigenetic reprogramming to attain pluripotency of the highest quality. While reprogramming of wt cells after pluripotency entry follows one pathway of sequential *Tet1* and *Oct4* gene activation, cells deficient of TET1 can presumably take an alternative route to reach induced pluripotency, albeit of an inferior status flawed with epigenetic defects. Recent efforts to coerce human ESC lines, which resemble primed-state murine EpiSCs, into naive-like states using alternative culture conditions have shown only a modest increase ^67,68^ or no detectable change ^4^ in *TET1* expression. Further improvement in reprogramming human ESCs and iPSCs to attain more features of naive pluripotency and genomic stability should therefore strategize at achieving full gene activation of *TET1*.

To our knowledge, our analysis provides unprecedented temporal resolution of global gene expression and DNA methylation changes in the final stages of iPSC generation, enabling dissection of nuanced differences in molecular signatures in the continuum of cell states during pluripotency acquisition. Our analysis of these late-intermediate stages contributes to a more complete molecular roadmap of reprogramming. Such dual reporter iPSC systems offer tractable models for further genomic analysis of pluripotency network rewiring, including the dynamics of histone modification, chromatin accessibility and transcription factor occupancy changes in the final phases of epigenetic reprogramming.

## Author Contributions

M.B. performed all experiments involving generation and characterization of transgenic mouse strains, embryos and MEF, cell culture, reprogramming colony time-course imaging and scoring, FACS and sorting, immunofluorescence staining, targeted bisulfite sequencing, telomerase assay, neural differentiation and all library preparations for RNA-seq and WGBS. X.L. performed RNA–seq analysis and comparative analysis with published databases. B.V. and B.T. contributed integrative RNA-seq and WGBS analysis and performed comparative WGBS analysis with published datasets. R.K. contributed expertise with mouse breeding and embryo dissection and assisted with time-course imaging. A.J. and V.P. contributed immunofluorescence staining and colony counting. J.X. contributed to reprogramming experiments and colony scoring. V.P. contributed expertise on reprogramming. C.V. provided funding and infrastructure support. M.B. wrote the methods, prepared all figures and contributed to the writing of the manuscript. K.P.K. conceived the study, directed the research and wrote the manuscript with assistance from all co-authors.

## Acknowledgements

We thank Rita San-Bento at Cambridge Epigenetix (Cambridge, U.K.) for technical guidance in oxidative WGBS library preparation. WGBS was performed at GenomeScan B.V. (Leiden, the Netherlands) with expert bioinformatics support provided by Thomas ChinAWoeng and Amrish Mahes. We thank Bernhard Payer and Konrad Hochedlinger for providing the transgenic OKSM mice. We thank Natalie De Geest for breeding OKSM mice, Lutgarde Serneels, Zhiyong Zhang and Liesbeth Vermeire (Mutamouse, KU Leuven) for the generation of Tg(*Tet1-*mCherry)B mice, Pier Andrée Penttila and Rob van Rossom (FACS Core, KU Leuven) for cell sorting expertise and Jan Cools for generous use of FACS MACSQuant^®^ VYB. This work was supported by the Fonds voor Wetenschappelijk Onderzoek (FWO) Research Foundation – Flanders Odysseus Program grants G.0C56.13N (K.P.K) and G0F7716N (V.P.), Research Project grant G.0632.13 (K.P.K), Ministerie van de Vlaamse Gemeenschap (K.P.K.) and the Marie Curie Career Integration Grant PCIG-GA-2012-321658 (K.P.K.), KU Leuven Internal Funds C14/16/077 (K.P.K. and V.P.) and SCIL program financing (C.V. K.P.K. and V.P). A.J. was supported by FWO PhD fellowship 1158318N; X.L. was supported by a PhD scholarship from the China Scholarship Council.

## Competing financial statements

The authors declare no competing interests.

## Methods

### Cell culture

Murine v6.5 embryonic stem cells (mESCs) (C57BL/6 × 129S4 F1 hybrid) and induced pluripotent stem cell (iPSC) lines were maintained on mitomycin C-inactivated primary mouse embryonic fibroblasts (feeders). The standard ESC culture medium consists of knockout DMEM (Cat.#10829-018, Thermo Fisher Scientific unless otherwise stated), 15% ES-qualified FBS (Cat.#10270-106) or KnockOut Serum Replacement (KSR) (Cat.#10828-028), 0.1 mM each of non-essential amino acids (Cat.#11440-035), 1 mM sodium pyruvate (Cat.#11360-039), 2 mM L-glutamine (Cat.#25030-024), 0.1 mM β-mercaptoethanol (Cat.#31350-010), 150 units penicillin/streptomycin (Cat.#15140), supplemented with in-house leukemia inhibitory factor (LIF) culture supernatant. Mouse embryonic fibroblasts (MEFs) and HEK293T cells were cultured in DMEM containing GlutaMAX (Cat.#61965-026), 10% FBS (Cat.#F7524, Sigma-Aldrich), 0.1 mM each of nonessential amino acids (Cat.#11440-035), 1 mM sodium pyruvate (Cat.#11360-039), 2 mM L-glutamine (Cat.#25030-024), 0.1 mM 2-mercaptoethanol (Cat.#31350-010) and 150 units penicillin/streptomycin (Cat.#15140). Serum-cultured ESCs were adapted to ground state conditions as previously described ^29^, by at least 5 passages in defined media referred as 2 inhibitors + LIF (2iL). The latter is composed of N2B27 basal media prepared from a 1:1 mixture of DMEM-F12 (Cat.#11320-074) and Neurobasal medium (Cat.#21103-049) supplemented with N2 (Cat.#17502-048), B27 (Cat.#17504-044), 1 mM L-glutamine, 0.1 mM 2-mercaptoethanol and 150 units penicillin-streptomycin. This basal media is further supplemented with 1 μM MEK inhibitor PD0325901 (Cat.#Axon 1386, Axon Medchem BV), 3 μM GSK3β inhibitor CHIR99021 (Cat.#Axon 1408, Axon Medchem BV) and 1000 U/ml ESGRO LIF (Cat.#ESG1107, Millipore). Tissue culture wells were pre-coated with 0.1% gelatine (Cat.#ES-006-B, Millipore), or 7.5 μg/ml poly L-ornithine (Cat.#P3655-50MG Sigma-Aldrich) and 5 μg/ml laminin (Cat.#L2020-1MG Sigma-Aldrich) for 1-2 hours at 37°C before seeding.

Primordial germ cell-like cells (PGCLCs) were generated as previously described ^30^. Briefly, murine ESCs were passaged for minimum five times in 2iL media. At day 0 they were seeded at 2-3 × 10^5^ per 10 cm^2^ and differentiated into epiblast-like cells (EpiLC) for 48 hours in N2B27 media supplemented with 1% KSR, 12 ng/ml bFGF (Cat.#100-18C, Peprotech) and 20 ng/ml Activin A (Cat.# 120-14E, Peprotech) on dishes pre-coated with 5 μg/10 cm^2^ fibronectin. At day 2, EpiLC were disaggregated with TrypLE Express (Cat.#12605010). 2 × 10^3^ cells were then seeded in each well of a 96-well lipidure plate (Cat.#AMS.51011610, Amsbio) and maintained in GK15 media composed by Glasgow’s MEM (GMEM) (Cat.# 11710035) 15% KSR, 0.1 mM each of non-essential amino acids, 1 mM sodium pyruvate, 2 mM L-glutamine, 0.1 mM β-mercaptoethanol, 150 units penicillin/streptomycin and 1000 U/ml ESGRO LIF. GK15 media was supplemented with 10 μg/ml BMP4 (Cat.#120-05ET, Peprotech), 10 μg/ml SCF (Cat.# 250-03-50μG, Peprotech), 25 μg/ml BMP8a (Cat.# 7540-BP-025, R&D), 50 μg/ml EGF (Cat.# 315-09, Peprotech). After 6 days, aggregates were collected and incubated for 10 minutes at 37°C with 0.25% trypsin-EDTA (Cat.# 25200056). Then they were filtered with a 70-μM cell strainer (Cat#734-0003, VWR) and stained with antibodies for cell sorting. From 2 × 10^3^ cells seeded at day 2, 1-2 × 10^4^ cells were collected from each aggregate at day 8.

Epiblast stem cells (EpiSCs) were derived from E6.5 post-implantation mouse embryos as previously described ^69^. The basal media used in this derivation protocol consist of 50% IMDM (Cat.# 21980-32), 50% Ham F12 Glutamax (Cat.#31765-027), 0.5% BSA (Cat.#EQBAH65-0100, Europa Bioproducts Ltd), 1% Lipid 100X (Cat.#11905-031), 450 μM monothioglycerol (Cat.#M6145, Sigma), 15 μg/ml insulin (Cat.#1376497, Roche), 7 μg/ml transferrin (Cat.# 652202, Roche) and 150 units penicillin/streptomycin (Cat.#15140). On the day of use, this was freshly supplemented with 20% KSR (Cat.#10828-028), 12 ng/ml bFGF (Cat.#100-18C, Peprotech) and 20 ng/ml Activin A (Cat.# 120-14E, Peprotech). After the first 3-4 passages, the cells were re-adapted to a maintenance media, consisting of N2B27 media supplemented with 12 ng/ml bFGF (Cat.#100-18C, Peprotech) and 20 ng/ml Activin A (Cat.# 120-14E, Peprotech) on mitomycin C-inactivated primary mouse embryonic fibroblasts ^70^.

EpiSC-to-ESC conversion was performed as previously described^34^. Briefly, the day before the transfection, 2-3 × 10^5^ single cells EpiSCs were seeded on 5 μg/10 cm^2^ fibronectin-coated 6-well plates in media containing (1 μl/ml) ROCK inhibitor (InSolution™ Y-276320, Cat.#688001-500UG, VWR Calbiochem). In the morning of transfection, media was replaced without ROCK inhibitor. Transfection of 4 μg Nr5a1 overexpressing plasmid (gift of Austin Smith) was performed in 500 μl OptiMEM (Cat.# 31985070) with 10 μl of Lipofectamine™ 2000 Transfection Reagent (Cat.# 11668019). On the day after, the transfection media was switched to 2iL and changed every day for 6/7 days until reset embryonic stem cells (rESC) colonies were ready to be picked.

Neural differentiation was performed as previously described^71^. Cells split at least two times on feeders in KSR media were adapted in feeder-free on gelatin for two passages. At day 0, 4 × 10^6^ cells were placed onto bacteriological Petri dishes (Cat. #633102, Greiner) in 15 ml MEF medium to form cellular aggregates. Media was changed every other day and starting from day 4, 5 μM of retinoic acid (Cat. #R2625, Sigma) was added to the media. At day 8 cellular aggregates were dissociated and 2-3 × 10^6^ cells were seeded on coverglasses in 12-well plates pre-coated with poly L-ornithine and laminin in N2 media composed by DMEM/F-12 (Cat.#31330-038) supplemented with N2 (Cat.#17502-048), 25 μg/ml insulin (Cat.#1376497, Roche), 50 μg/ml 7.5% BSA and 2 mM L-glutamine (Cat.#25030-024). At day 8+2 media was replaced with B27 complete media composed by DMEM/F-12 (Cat. #31330-038) supplemented with B27 (Cat.#17504-044), and 2 mM L-glutamine; refreshed every other day. Cells were fixed at day 8+6 for immunofluorescent staining.

### Mouse strains

**Tg(Col1a-OKSM)**: B6;129S4-Col1a1tm1 (tetO-*Pou5f1*,-Klf4,-Sox2,-Myc)Hoch/J^37^. This strain (Jackson stock 011001) was obtained from Bernhard Payer and Konrad Hochedlinger.

**Tg(*Pou5f1*-EGFP)** expressing “naive” *Oct4-*GFP: B6;CBA-Tg(*Pou5f1*-EGFP)2Mnn/J ^31^. The transgene contains an Enhanced Green Fluorescent Protein (EGFP) gene under the control of the promoter and distal enhancer of the POU domain, class 5, transcription factor 1 (*Pou5fl*). The TgOG2 transgenic mice line, obtained from Jackson (stock 4654), was generated by pro-nuclei injection of (CBA/CaJ×C57BL/6J)F2 zygotes and has been maintained by homozygote × homozygote breeding.

***Pou5f1***^**tm2Jae/J**^ expressing “total” *Oct4-*GFP: B6;129S4-*Pou5f1*^tm2Jae/J^ ^32^. A targeting vector was designed to insert an “IRES-EGFP-floxed NEO” cassette (containing an internal ribosomal entry site (IRES), EGFP sequence, and a loxP-flanked neo) between the stop codon and endogenous polyA signal of the targeted gene. This strain was obtained from Jackson (stock 8214) at N2F? and backcrossed in-house to C57BL/6J for another two generations prior to interbreeding with B6-Tet1^tm1.1Koh^.

**Tg(*Tet1-*mCherry)B** expressing “naive” *Tet1-*mCherry: B6;129S-Tg(*Tet1-*mCherry)B ^25^. The transgene contains mCherry-2A-puromycin resistance ORF cassette under the control of the distal promoter (6-kb genomic region B) of the gene for *Tet1*. Both clonal transgenic lines (B37 and B43) were backcrossed to C57BL/6J for 3 generations and thereafter maintained by filial generation matings. N3F? mice were subsequently bred to TgOG2 or *Pou5f1*^tm2Jae/J^.

***Tet1***^**tm1Koh**^ ^27^. A target vector was designed to insert an “fl-NLS-lacZ-fl-mCherry-FRT-neo-FRT” cassette (containing ATG start codon and nuclear localization signal sequence 5’ of lacZ reporter gene with 3’ BGH polyA, flanked by 2 LoxP sites to allow reporter exchange to a downstream ATG-mCherry-pA cassette by Cre recombination, followed by a FRT-flanked neomycin selection cassette) in-frame at the ATG start codon in exon 2 of *Tet1*. The targeting construct was electroporated into C57BL/6J mouse embryonic stem (ES) cells and selected in G418. The transgenic KI mouse strain was created by blastocyst injection of one correctly targeted clone. The strain has been bred to Flp-deleter mice to remove the neo cassette, resulting in one remaining FRT site downstream of the fl-NLS-lacZ-fl-mCherry cassette. The final modification is designated as *Tet1-*fl-lacZ-fl-mCherry.

***Tet1***^**tm1.1Koh**^ expressing “total” *Tet1-*mCherry (B6-*Tet1*^tm1.1Koh^). The B6-*Tet1*^tm1Koh^ line was crossed to B6-Cre-deleter mice to remove the NLS-lacZ cassette, resulting in one loxP site at the 5’end of ATG-mCherry-pA. The final modification is designated as *Tet1-*fl-mCherry.

Dual reporter strains were interbred to double homozygosity whenever possible, with the exception of the *Tet1*^tm1^ or *Tet1*^tm1.1^ alleles which were maintained as heterozygotes. Time pregnancies were set up between double reporter females and homozygous OKSM males to collect mouse embryonic fibroblasts. All experimental procedures on mice have been reviewed and approved by the KU Leuven Ethical Committee for Animal Experimentation (P101/2016) in compliance with European Directive 2010/63/EU.

### Characterization of Tg(*Tet1-*mCherry)B ESC lines and creation of transgenic mice

v6.5 ESCs containing the Tg(*Tet1-*mCherry)B reporter transgene were generated as described in Sohni *et al.*, 2015. Southern blot analysis of Tg(*Tet1-*mCherry)B clones was performed to select two clones that contain intact cassettes at single random integration sites. Genomic DNA (gDNA) was purified from transgenic mESCs using the PureLink Genomic DNA Mini Kit (Cat# K1820-02, Invitrogen) according to manufacturer’s instructions. Approximately 10 μg of gDNA was digested with StuI-HF restriction enzyme (Cat# R0187M, NEB). Genomic fragments were then resolved on a 0.7% agarose gel for 4/5 hr at 140V in TBE buffer and transferred overnight from the gel via capillary action to a Zeta-probe membrane (Cat#162-0165, Biorad). Probe 4 (see Supplementary Figure 1a) was amplified by PCR and labeled with [α-32P]dCTP using TaKaRa Bio Ladderman Labeling kit (Cat. #6046, Westburg) according to the manufacturer’s instructions. Primer sequences are listed in Table S1. Hybridization, washes and detection were performed following a standard protocol as recommended by ExpressHyb (Cat#636831, Westburg). Based on our Southern strategy, the endogenous region is expected to be detected as a 10 kb band, whereas the incorporation of an intact transgene reporter elsewhere should be detected as an additional 6 kb band. By these criteria, clones 37 and 43 were selected for further characterization.

To map the (*Tet1-*mCherry)B transgene insertion sites precisely, genomic DNA samples from clones 37 and 43 were sequenced by targeted locus amplification ^28^ (Cergentis B.V, Utecht, the Netherlands). This method validated single site integration of the transgene at chromosome 10 in two different locations without disruption of the endogenous genes (Figure S1D-G). These two ESC clones were injected into CD1 host blastocysts to generate chimeric offspring at the VIB-KUL InfraMouse facility. Germline transmission was confirmed by coat color and genotyping, following which transgenic F1 generation was backcrossed with C57BL/6 mice for three generations.

### Genotyping

Mouse tail tip or 13.5-14.5 dpc embryo heads were digested in 500 μl lysis buffer (100 mM Tris-Cl pH 8.5, 200 mM NaCl, 5 mM EDTA, 0.2% SDS) supplemented with 200 μg/ml Proteinase K (Cat.# 3115879001, Sigma-Aldrich) at 55°C overnight in an orbital shaking heat block. After clarifying the tissue lysate by centrifugation at ≥8000 g for 10 minutes, the supernatant was mixed with an equal volume of isopropanol to precipitate genomic DNA. The pellet was dissolved at 55°C for 30 min in 300 μl milliQ water. 1 μl of DNA (80-100ng/μl) was used for genotyping by PCR, using GoTaq G2 Flexi DNA Polymerase (Cat.# M7805, Promega) according to the manufacturer’s instructions. Primer sequences are listed in Table S1.

### Embryo isolation and mouse embryonic fibroblast (MEFs) derivation

To collect pre-implantation embryos, 4- to 5-week old female mice were super-ovulated by intraperitoneal injection of 5 IU pregnant mare serum gonadotropin (Cat.#G4877 Sigma-Aldrich) followed by 7.5 IU human chorionic gonadotropin (Cat.# CG5-1VL, Sigma-Aldrich) 46-48 h later, and then mated with stud males. At 3.5 days post-coitum, blastocysts were collected by flushing the uterus with M2 media (Cat.# MR-015-D, Millipore). Embryos were imaged using the Nikon Ti2 inverted microscope imaging system.

Post-implantation embryos (E6.5) were isolated from decidua dissected from the uterine lining of naturally mated mice. Following removal of the Reichert’s membrane using forceps, the intact embryos were imaged using Nikon Ti2 inverted microscope imaging system. To derive EpiSCs, 6.5 dpc embyos were placed in droplets of 2.5% pancreatin/0.5% trypsin in Ca^2+^/Mg^2+−^ free Tyrode Ringer’s saline solution for 5-10 minutes until the visceral endoderm started to detach. Embryos were transferred to a new 60 mm dish containing 5 mL of sterile MEF medium where visceral endoderm was removed by mechanical dissection. Subsequently, embryonic and extra embryonic tissues were separated. The epiblast was washed once in EpiSC medium and moved to a well of 24-well plate of feeders containing EpiSC medium.

MEFs were isolated from 13.5 or 14.5 dpc mouse embryos. Briefly, the heads, visceral organs and gonads were removed under a stereomicroscope. Head tissues were used for genotyping and gonads were examined for gender identification. Individual carcasses were mechanically disaggregated with blades and incubated for 10 minutes at 37°C with 0.25% trypsin-EDTA (Cat.# 25200056, Thermo Fisher). The homogenate was passed twice through a 19G syringe needle (Cat.# 613-3938, VWR) before plating in T150 flasks and incubated at 37°C for 48-72 hours. Cell lines were frozen at the first passage.

### Lentiviral production

The day before transfection, HEK293T cells were passaged from 90-100% confluent 10-cm dishes into 15-cm dishes in MEF media. On the day of transfection, plasmid DNA was prepared in OptiMEM (Cat.# 31985070, Thermo Fisher). To produce virus for STEMCCA expression, plasmids were transfected in the ratio of 30 μg pHAGE-tetO-STEMCCA: 1.5 μg tat: 1.5 μg rev: 1.5 gag/pol: 3 μg vsv-g ^35^. To generate viruses for rtTA and shRNA, plasmid transfection ratios were respectively 12 μg FGΔGW-M2-rtTA: 9 μg △8.9: 6 μg vsv-g ^8^ and 7.74 μg pLKO.1 shRNA plasmid: 5.8 μg psPAX2: 1.8 μg pMD2.G. Trans-IT (Cat.# MIR2700, Mirus) was used at 3(μl) volumes per μg of DNA according to manufacturer’s instruction. The day after transfection the media was changed and supernatants were collected from the transfected cells starting at 48 hours after transfection once every 12 hours (4-5 collections), filtered through 0.45-μm (Cat.# VWR 514-4101) and pooled. Viruses were concentrated by spinning for 2 hours at 3500 rpm, 4°C in Vivaspin20 columns (Cat.#VS2032,Sartorius) and stored at –80°C.

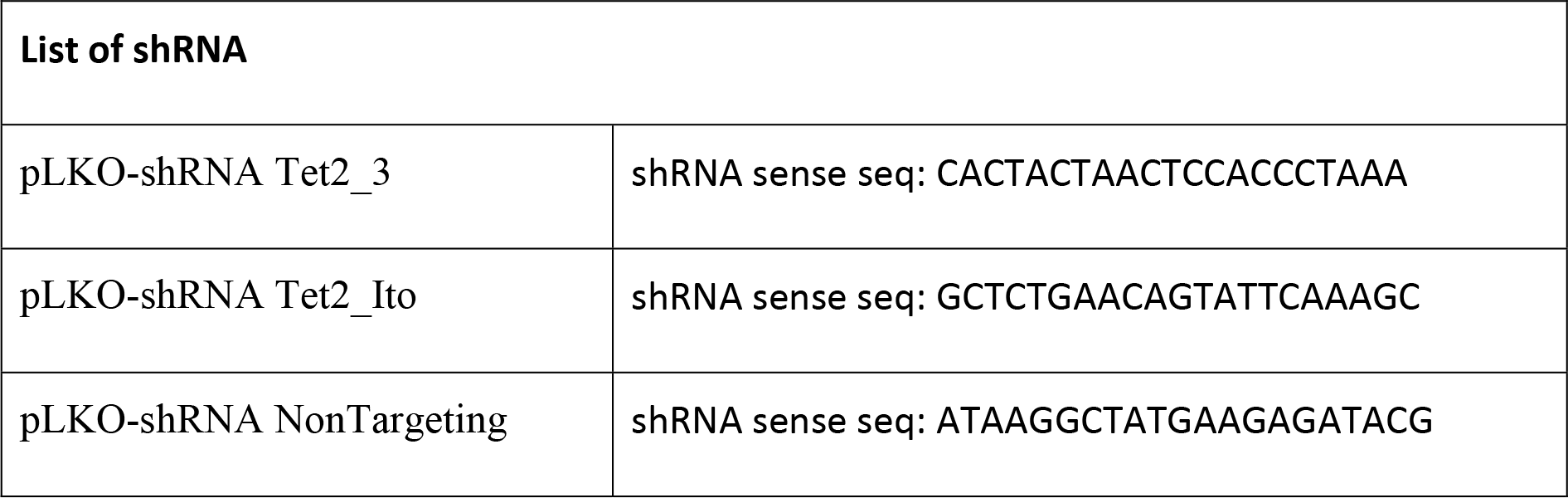

Titering of viral supernatant was performed by seeding 5 × 10^4^ MEFs or ESCs (for shRNA) in each well of a 12-well plate the day before transfection. Different amounts of viral supernatant in roughly two-fold increments (10μl to 200μl pHAGE-tetO-STEMCCA, 10μl to 100μl FGΔGW-M2-rtTA and 2μl to 32μl of shRNA) were added to the cells dropwise on 0.5-ml of fresh culture supernatant containing 5 μg/ml polybrene (Cat.# TR-1003-G, Millipore). On the following day, 1 μg/ml doxycycline (Cat.#D9891-1G, Sigma-Aldrich) was added to the media. After 48h cells were fixed in 4% paraformaldehyde/PBS and stained with mouse anti-Oct-3/4 (C-10) monoclonal antibody (1:250, Cat.#sc-5279, Santa Cruz). Cells were imaged using Zeiss Axiovert 40 CFL microscope and Oct4 positive cells of all DAPI-stained cells were scored with ImageJ software to determine the best titer. Cells infected with shRNA were kept for 48h after infection, then harvested for RNA and titrated by qPCR for the target gene.

### MEFs to iPSCs reprogramming

MEFs containing dual reporters without transgenic OKSM (including *Tet1-*KO paired with wt) were infected with pooled concentrates of dox-inducible pHAGE-tetO-STEMCCA and reverse tetracycline transactivator (FGΔGW-M2-rtTA)-expressing lentivirus. MEFs of passage 3 were seeded at 5 × 10^4^ cells per well of a 12-well plate and infected the next day with lentivirus added dropwise onto 0.5-ml of fresh culture supernatant containing 5 μg/ml polybrene (Cat.# TR-1003-G, Millipore). On the following day (day 0), culture supernatant was switched from MEF to standard ESC medium containing 1 μg/ml doxycycline (Cat.#D9891-1G, Sigma-Aldrich) with or without 50 μg/ml ascorbic acid (Cat.# A4544, Sigma-Aldrich) until day 10. Subsequently, ESC media with or without 50 μg/ml ascorbic acid was refreshed every day in the absence of doxycycline. iPSC colonies were picked at day 16 and expanded to establish iPSC lines. All experiments where reprogramming intermediates were fixed for immunofluorescence staining were performed as described above, in serum containing ascorbic acid.

In experiments involving Tet2 shRNA knockdown, MEFs were first infected with pHAGE-tetO-STEMCCA and rtTA as described above and treated with doxycycline at day 0. At day2, the MEFs were further transduced with shRNA lentiviral supernatant and selected with 1.5 μg/ml puromycin for 2-3 days. On day 4, cells were passaged 1:4 on feeders cells; on day 5 the media was changed into KSR mESC media.

MEFs containing Tg(Col1a-OKSM) were seeded at 2 × 10^4^ cells per well of a 12-well plate and infected with FGΔGW-M2-rtTA lentivirus in media containing 5 μg/ml polybrene the day after. On day 0, media was replaced with ESC media containing 2 μg/ml of doxycycline with or without ascorbic acid (50 μg/ml). Experiments performed in KSR and ascorbic acid media, which showed faster reprogramming kinetics, were started as previously described; then on day 2, cells were passaged 1:4 on feeders cells and on day 4 the media was changed into KSR ESC media in which KSR substitutes for FBS.

Reprogramming efficiencies were assessed by alkaline phosphatase staining of iPSC colonies formed by days 14-16. Cells were washed once with PBS, fixed with 4% paraformaldehyde for 5 minutes and stained using the Vector Red Substrate kit (Cat#SK5100, Vector Laboratories) according to kit instructions. After staining, cells were washed twice with PBS for imaging and manual colony counting.

### Fluorescent images

Images of reprogramming cultures were taken daily by using InCell analyser 2000 (GE Healthcare Life Sciences) (data Figure 2 and S2) or Nikon Ti2 automated fluorescent image scanner (data Figure 3 and S3). In either systems, pre-specified 9 × 9 fields per well centered on a 6-well format were captured using an automated x-y motorized stage. Bright field, mCherry and GFP were imaged every day from day 7 to day 16 for each defined field. Image fields containing iPSC colonies at the terminal time point were retrospectively scored along the time course series for the sequential events of fluorescence detection. Images acquired per well on the Nikon Ti2 were electronically stitched using NIS-Element software (Nikon) for full-well image reconstitution. Scoring was performed manually on ImageJ. PGCLCs, EpiSCs and rESCs cell cultures from Figure 1 were imaged with Nikon Ti2 automated fluorescent microscope.

### Fluorescence activated cell sorting (FACS)

Transgenic ESC lines and reprogramming intermediates were trypsinized, filtered and analysed by flow cytometry on a MACSQuant^®^ VYB (Miltenyi Biotec) for all data acquisition involving mCherry detection. Tet1KO and WT reprogramming intermediates were incubated 30 min on ice with Phycoerythrin (PE)-conjugated Mouse anti SSEA1 (1:100, Cat.#FAB2155P-025, R&D) and analysed on CANTO I (BD Biosciences). PGCLCs at day8 were sorted using FITC Mouse anti-SSEA1 (1:20, Cat.#560127, BD Biosciences) and BV421 Hamster Anti-Mouse CD61 (1:50, Cat.#562917, BD Biosciences). All cell sorting experiments were performed with BDFACS Aria III (BD Biosciences). FACS data were plotted by using FlowJo software.

### Magnetic cell isolation

Reprogramming intermediates were labelled with Anti-SSEA-1 (CD15) MicroBeads (Cat.#130-094-530, Miltenyi Biotec) and magnetically isolated according to the manufacturer’s instructions. The labelled cells were loaded onto a MACS LS Column (Cat# 130-042-401, Miltenyi Biotec), which is placed in the magnetic field of a MACS Separator. SSEA1+ cells were retained within the column while unlabelled cells run through. After removing the column from the magnetic field, SSEA1+ cells were eluted.

### Immunostaining

All IF experiments in Figures 2I-J, 2O-P, S2F, 3E-J and S3G-J were performed by lentiviral OKSM transduction and reprogramming in serum + AA conditions using cells seeded on gelatinized glass coverslips in 12-well plates. At defined time points, cells were fixed in 4% paraformaldehyde for 10 min at RT and permeabilized with 0.5% Triton-X100 in PBS for 5-10 min. Fixed cells were then incubated in PBS containing 5% donkey serum, 0.02% Tween20 and 0.02% fish skin gelatin with primary antibodies for DPPA4 1:250 (Cat.#AF3730,R&D), mCherry 1:1000 (Cat.#ab167453,Abcam), TET1 1:200 (gift of Heinrich Leonhardt ^40^), GFP 1:500 (Nacalai Tesque Cat# 04404-84) or NANOG 1:200 (Cat.#ab80892,Abcam) overnight at 4°C in blocking solution. Following incubation with the appropriate fluorophore-labeled secondary antibodies in blocking solution for 40 minutes in the dark and counter-staining with DAPI (Cat.# D9542-5MG, Sigma-Aldrich) with intervening washes with 0.2% Tween in PBS (PBST), slides were mounted in ProLong Gold antifade reagent (Cat.#ab P36930,Thermo Fisher) and imaged using a Zeiss AxioImager Z1 fluorescent microscope. Colonies with minimum four touching cells were scored. At early time points (e.g. day 6-8) the full coverslip was scored while at late time points (e.g. day 10-14) ~100 colonies were scored.

Cells at day 8+6 of neural differentiation were stained with 1:500 TUJ-1/β3TUB antibody (Cat.# 302302,Synaptic System) and counter-stained with DAPI (Cat.# D9542-5MG, Sigma-Aldrich).

### Quantitative reverse transcription-PCR (Q-PCR)

Total RNA was extracted by using the RNeasy mini-kit (Cat#74004, Qiagen) according to the manufacturer’s instructions. 0.5-1.0 μg RNA was converted to cDNA using SuperScript III (Cat#11752-050, Thermo Fisher). Q-PCR was carried out using cDNA input used at 1:100 in SYBR green PCR master mix (Cat#11733-046,Thermo Fisher) on a StepOnePlus (96-well) or ViiA7 real-time (384-well) PCR system (Life Technologies). In copy number measurements, absolute standard curves were generated by 10-fold serial dilutions of plasmid clones containing target sequences, starting from 10^5^ or 10^6^ copies estimated based on mass concentration. Transcript copy numbers were determined by calibration to standard curves and normalized to *Gapdh* gene copy numbers. Relative expression was calculated using ΔΔCt method, using Gapdh expression for normalization and a reference control sample. Primer sequences are listed in Table S1.

### RNA sequencing

1×10^5^ to 3×10^6^ cells in each reprogramming intermediate fraction were sorted from three independent lines (starting from transgenic MEFs isolated from individual embryos) as biological replicates. RNA was purified using TRIzol (Cat.#15596026,Invitrogen) and RNeasy mini kit (Cat#74004, Qiagen). Purified total RNA was quantified by using the Agilent RNA 6000 Nano kit (Cat# 5067-1511, Agilent) on an Agilent 2100 Bioanalyzer (Agilent). 300 ng was used for library preparation using the KAPA stranded mRNA-seq kit (Cat#07692193001, Roche) according to the manufacturer’s instructions (V5.17). KAPA single-indexed adapters (Cat#08005699001, Roche) were added to the A-tailed cDNA for library amplification with 14 cycles of PCR. The final library was purified by using Agencourt AMPure XP beads (Cat#A63881, Beckman Coulter) to size select fragments of 200-300 bp. Library size was validated by using the High Sensitivity DNA Analysis kit (Cat# 5067-4626, Agilent) on an Agilent 2100 Bioanalyzer (Agilent). Library DNA concentration was measured by using the Qubit HS dsDNA Quant kit (Cat#Q32851, Thermo Fisher). Each library was diluted to 4 nM and pooled for single-end 75-bp sequencing on an Illumina NextSeq500 for 15-24 million reads per sample.

Index-trimmed single-end 75-bp reads were aligned to the mouse reference genome (mm10) by using Hisat2 (v2.1.0) to generate BAM files with 75-90% unique mapping reads. Gene-level read count matrix was summarized from mapped reads by using featureCounts (v1.5.3; http://bioinf.wehi.edu.au/featureCounts/). Subsequently, the count matrix was imported into the R Bioconductor DESeq2 package (v1.18.1) for differential gene expression analysis with false discovery rate (FDR) < 0.05 and different log2 fold change thresholds (magnitude > 1, 1.5 or 2). Differentially expressed genes were collated to generate heat-map clusters by using R package pheatmap (v1.0.8) with scaling and clustering only the rows. Principle component analysis (PCA) was performed using plotPCA function from DESeq2 package with unsupervised transformed counts. R build-in function hclust() was used to hierarchically cluster differential expression genes across groups. Enriched gene ontology (GO) terms were generated using the R Bioconductor clusterProfiler package (v3.6.0;) with default parameters (adjusted *P* value < 0.05 and |log_2_ fold-change (FC)| > 1). Normalized gene expression values (Transcripts Per Kilobase Million, TPM) were quantified using Salmon (v0.8.1, https://combine-lab.github.io/salmon/) with default parameters using the total gene annotation of the ENSEMBL genome assembly GRCm38 transcript release 85 version.

For combined analysis with the RNA-seq dataset from Knaupp et al. (2017), GRCm38 transcripts release 85 version was filtered by excluding transcripts on sex chromosomes, the mitochondria genome, all histone-coding transcripts and all NCR’s (non-coding RNAs) to remove the bias led by different RNA-seq library prep methods. Then both RNA-seq datasets were re-quantified with this modified reference using Salmon to obtain TPM values. To generate the combined PCA plot, the TPM values of all differential expression genes of 5 comparisons from Bartoccetti’s RNA-seq data and 5 comparisons from Knaupp’s RNA-seq data were used.

### Bisulfite sequencing

Genomic DNA was purified by using PureLink™ Genomic DNA Mini Kit (Cat# K182001, Thermo Fisher). 400 ng of DNA was bisulfite treated by using the EpiTect bisulfite conversion kit according to the manufacturer’s protocol (Cat#59104, Qiagen). PCR products were generated by using Hifi Taq polymerase (Cat#11304-011: Thermo Fisher) and subcloned into the pGEM-T Easy vector (Cat#A1360, Promega). At least 10 clones were picked for Sanger sequencing (GATC biotech). The CpG methylation status of the sequences was analyzed using QUMA^72^. Primer sequences are listed in Table S1.

### Whole Genome Bisulfite sequencing (WGBS)

1×10^5^ to 3×10^6^ cells in each sorted fraction of reprogramming intermediates were sorted in biological replicates. DNA was isolated by using DNeasy Blood & Tissue Kit (Cat# 69504, Qiagen) according to the manufacturer’s instructions. 200 ng of DNA was subjected to oxidative bisulfite sequencing (OxBS) by using TrueMethyl Whole Genome kit v3.1 (CEGX) according to the oxBS mode only pipeline. λ-DNA was added to the mix as 0.5% of total DNA amount for calculation of BS conversion rates. Libraries were sequenced by 150-bp paired-end reads on an Illumina HiSeq 4000 platform (GenomeScan, NL) to generate 45 Gb data per sample. Shallow run was performed on 10 samples (~3-fold genome coverage) and eight of them were sequenced in full depth (total ~20-fold coverage). Bisulfite conversion rates ranged from 98.3-99.3% and oxidation conversion rates from 93.0-95.5% based on internal spike-in controls of the TrueMethyl kit. Quality trimming of reads included removal of presumed adapter sequences using the in-house tool AdapterTrim (v2), low complexity trimming according to TrueMethyl kit instructions to eliminate most Adaptase tails, and filtering using QualityTrim (v2) to remove bases with phred scores below Q30 and reads shorter than 36 bp (both pairs in paired-end reads). Resulting quality-trimmed reads averaged 129.4 ± 2.3 bp with phred scores 39.1 ± 0.2 (mean ± SD) across all samples. Read pair duplicates (8.0-15.9%) were removed using the deduplication program provided by Bismark, resulting in final read pairs in the range of 93.3-101.9 million. Reads were aligned to UCSC mm10 reference sequence using Bismark (v0.19.0) with default settings, yielding mapping rates of 61.5-67.8% across all samples. Methylation calls at all covered Cs were extracted with Bismark extraction to calculate global methylation percentages. Methylation levels at individual Cs were calculated by Bismark using thresholds set with a minimum coverage of 10 and a maximum coverage with a percentile of 99, as recommended by the Methylkit package. WGBS data (SRR5870485 and SRR5870484) from Schwarz et al. (2018) were re-mapped to the mouse genome (mm10), and methylation calls were extracted using Bismark (v0.19.0) in a similar fashion ^50^.

Correlation between samples were calculated using the default setting with the MethylKit (v1.4.1). Each single CpG called was annotated with MethylKit and was assigned to one of the following categories; Intron, exon, IAP, LINE, maternal, paternal, promotor, CpG islands (CGI), CGI shores, 5’UTR, 3’UTR and intergenic. The USCS mm10 bed file was used as reference for this process. Probes overlapping with CGIs in each category were reclassified as separate CGI classes within promoters and outside (non-promoter). The generated results were visualised as boxplots (Figure 5E). For the annotation of the maternal and paternal germline imprint control regions, the minimum threshold coverage was changed to 5. Annotations to define maternally and paternally imprinted regions are based on Wang et al. (2014) (ref 49). Methylation levels for these regions were calculated as the number of methylated CpGs over all methylated and un-methylated CpGs meeting the threshold. PCA shown in Figure S6L was based on methylation levels for covered single CpG sites).

Probes for 1-kb tiles and promotors of DEGs were generated using SeqMonk (v1.44.0) and subsequently further processed using a custom R script (R version 3.5.2). Promotors were defined as the region starting at 2 kb upstream and ending 500 bp downstream of the most upstream TSS annotated on ENSEMBL version 90. The methylation level for each region was calculated as the sum of all methylated CpG measurements, divided by all CpGs measurements (methylated and unmethylated). Tiles and promoters were excluded if fewer than 20 CpG methylation measurements were made, or if more than the 99^th^ percentile of CpG methylation measurements were made. This resulted in a final selection of 62.1% of all tiles and 91.1% of all promotors for further downstream analysis.

Heatmaps for methylation were made using the complex Heatmap package (v1.20.0)^73^ using no scaling and only row-clustering based on the Euclidian distance (Figure 5G, Figure S5I). Cluster analysis of 1-kb tiles was done using the k-means clustering method (stats R-package, v3.5.4) with the MacQueen algorithm, a max iteration of 1000, 10 random sets, and k = 10. Based on the observed demethylation kinetics of the clusters, the 1-kb tiles were categorized into 2 groups: early and late. PCA using 1-kb tiles (Figure 6E) was performed using the R prcomp function (stats R-package, v3.5.2). DMRs of wt and KO iPSCs were calculated using the tool Defiant^74^ with default parameters, using a threshold set on a minimum of 10x coverage and 25% methylation percentage difference. Genomic annotation of DMRs was done using the ChipSeeker R-package (v1.18.0)^75^. Functional annotation of the different groups was done with Bedtools annotate (Bedtools2, v2.25.0) and an ESC-specific ChromHMM genome annotation as a reference ^76^. DMRs were intersected with regions where there was also a 25% difference between iPSCs and MEFs that fell into the early and late demethylation groups using Bedtools intersect (Bedtools2, v2.25.0) and used as inputs in GREAT (v3.0.0)^77^ for GO analysis using default parameters (FDR < 0.05 and observations > 5 in both binomial and hypergeometric tests).

### Flow cytometry-fluorescence in situ hybridization (Flow-FISH) assay

A fluorescein-conjugated peptide nucleic acid (PNA) telomere detection kit for flow cytometry (Cat.#K5327, Dako) was used following the manufacturer’s protocol, except that fluorescein-conjugated PNA was substituted with an alternative probe. Briefly, 4×10^6^ EpiLC were harvested and separated in 4 tubes. Two tubes were hybridized overnight with 0.5μg/ml of Cy5-labeled G-rich telomere probe (Cat.#PN-TG055-005, Eurogentec) and two tubes were used as unlabeled control. After two washes, cells were stained for more than 3-4 hours with a solution containing propidium iodide provided in the kit and analyzed by FACS.

