## Supplemental Figures and Table for "Temporal resolution of global gene expression and DNA methylation changes in the final phases of reprogramming towards induced pluripotency"

### Figure S1

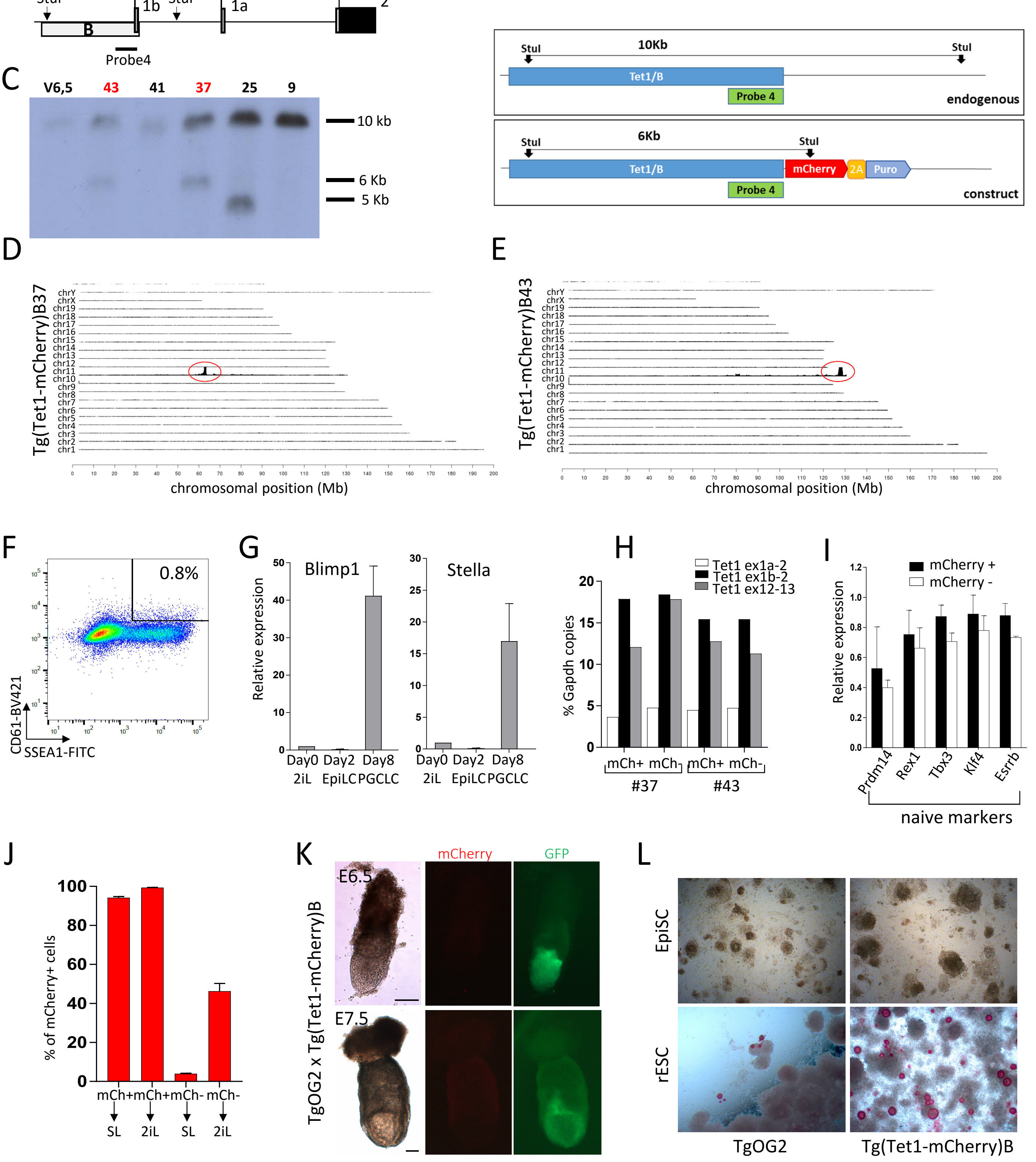
A
Tet1
StuI
1b
StuI
1a
ATG
2
B
Probe4
B
10Kb
StuI
StuI
Tet1/B
Probe 4
endogenous
6Kb
StuI
StuI
Tet1/B
mCherry
2A
Puro
Probe 4
construct
C
V6,5
43
41
37
25
9
10 kb
6 Kb
5 Kb
D
Tg(Tet1-mCherry)B37
chromosomal position (Mb)
E
Tg(Tet1-mCherry)B43
chromosomal position (Mb)
F
0.8%
CD61-BV421
SSEA1-FITC
G
Blimp1
Stella
Relative expression
Day0 2iL
Day2 EpiLC
Day8 PGCLC
H
Tet1 ex1a-2
Tet1 ex1b-2
Tet1 ex12-13
% Gapdh copies
mCh+ mCh-
#37
#43
I
mCherry +
mCherry -
Relative expression
Prdm14
Rex1
Tbx3
Klf4
Esrrb
naive markers
J
% of mCherry+ cells
mCh+ mCh+ mCh- mCh-
SL 2iL SL 2iL
K
TgOG2 x Tg(Tet1-mCherry)B
E6.5
E7.5
mCherry
GFP
L
EpiSC
rESC
TgOG2
Tg(Tet1-mCherry)B

**Figure S1**. **Related to Figure 1. Characterization of transgenic reporter strains.** (A) The 5′ genomic region of *Tet1* showing promoter region B. StuI restriction sites are indicated and the black bar marks the site detected by Southern probe 4. (B) Southern blot (SB) strategy to validate the integrity of Tg(*Tet1*-mCherry)B cassette electroporated in ESCs. (C) SB analysis of genomic DNA harvested from wild type (v6.5) and transgenic ESC clones. The endogenous region is detected as a 10-kb band, while an intact transgene reporter is detected as an additional 6-kb band (in clones 37 and 43). (D-E) Genome-wide coverage plot by targeted locus amplification (TLA) sequencing confirms a single transgenic insertion in both clones 37 (D) and 43 (E) on chromosome 10. (F) Flow cytometry plot of CD61+/SSEA1+ PGCLCs sorted at day 8. Gating indicates cell fraction sorted for analysis. (G) Relative expression of *Blimp1* and *Stella* in the sorted PGCLCs. Expression values are shown as means ± SEM (n=3 independent sorts). (H) Expression of different *Tet1* 5' transcript isoforms and 3' coding sequence (exons 12-13) in sorted mCherry-negative and -positive Tg(*Tet1*-mCherry)B ESCs from clones 43 and 37. (I) Relative expression of naive pluripotency marker genes in mCherry-negative and -positive transgenic ESCs. Expression values are means ± SEM (n=3; n=2 for clone 37 and n=1 for clone 43). (J) Percentage of sorted mCherry-negative and -positive cells passaged >7 times in SL or 2iL media. Representative FACS plot is shown in Figure 1F. Data are represented as means ± SEM (n=5). (K) Bright-field (left) and fluorescence images of post-implantation E6.5 and E7.5 TgOG2;Tg(*Tet1*-mCherry)B dual reporter embryos. Scale bars represent 100 μm. (L) Bright field images of alkaline phosphatase (AP) stained EpiSCs and rESCs of Tg(*Tet1*-mCherry)B and Tg(*Pou5f1*-EGFP)Mnn single reporter lines.

Figure S2

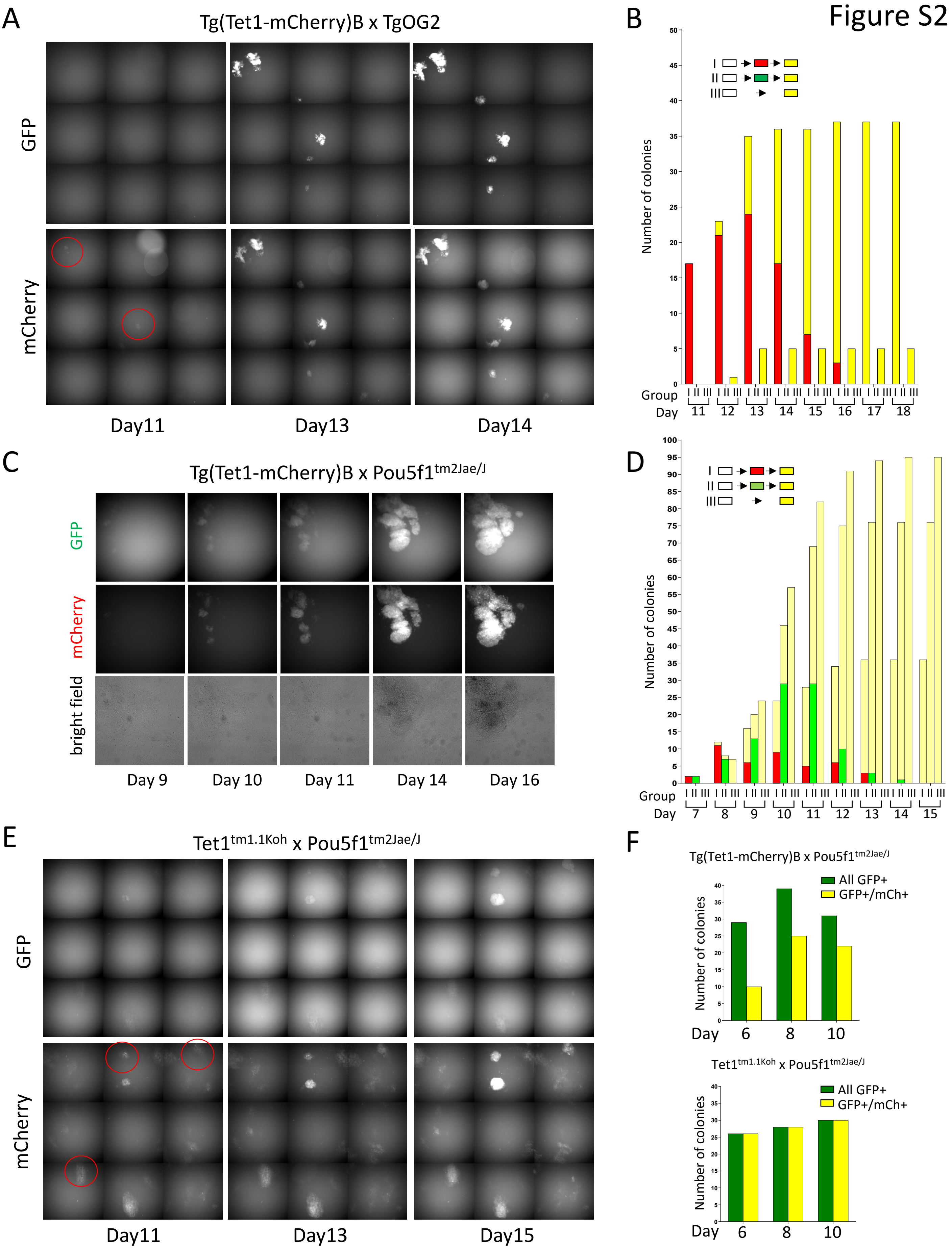
A
Tg(Tet1-mCherry)B x TgOG2
GFP
mCherry
Day11
Day13
Day14
B
Number of colonies
Group
Day
11 12 13 14 15 16 17 18
I
II
III
C
Tg(Tet1-mCherry)B x Pou5f1tm2Jae/J
GFP
mCherry
bright field
Day 9
Day 10
Day 11
Day 14
Day 16
D
Number of colonies
Group
Day
7 8 9 10 11 12 13 14 15
E
Tet1tm1.1Koh x Pou5f1tm2Jae/J
GFP
mCherry
Day11
Day13
Day15
F
Tg(Tet1-mCherry)B x Pou5f1tm2Jae/J
All GFP+
GFP+/mCh+
Number of colonies
Day
6
8
10
Tet1tm1.1Koh x Pou5f1tm2Jae/J
All GFP+
GFP+/mCh+
Number of colonies
Day
6
8
10

**Figure S2. Related to Figure 2. Kinetics of *Tet1* and *Oct4* gene activation during reprogramming.** (A-B) Replicate experiments to score dual reporter fluorescence activation of Tg(*Tet1*-mCherry)B; Tg(*Pou5f1*-EGFP)Mnn during reprogramming in serum media. (A) Stitched fluorescent images of selected fields in a representative time-course experiment. Red circles identify colonies that are mCherry positive and still GFP negative. (B) Daily colony counting of an independent experiment. (C-D) Replicate experiments to score dual fluorescence activation of Tg(*Tet1*-mCherry)B; *Pou5f1*$^{tm2Jae/J}$ during reprogramming in serum media. (C) Representative fluorescence images of a single colony imaged over time. (D) Daily colony counting pooled from 3 independent experiments (2 experiments using cell lines harboring Tg(*Tet1*-mCherry)B37 on N2 backcross to B6 and one using a cell line harboring Tg(*Tet1*-mCherry)B43 on CD1 background). (E) Stitched fluorescent images of a *Tet1*$^{tm1.1Koh}$; *Pou5f1*$^{tm2Jae/J}$ dual reporter line imaged at different time points in serum media. Red circles identify colonies that are mCherry positive and still GFP negative. (F) Number of colonies scored from immunofluorescence staining of mCherry and GFP in both Tg(*Tet1*-mCherry)B *Pou5f1*$^{tm2Jae/J}$ and *Tet1*$^{tm1.1Koh}$; *Pou5f1*$^{tm2Jae/J}$ dual reporter lines at different time points in serum+AA media. Green bars indicate total GFP+ colonies counted; yellow bars indicate GFP+ colonies that were also mCherry+.

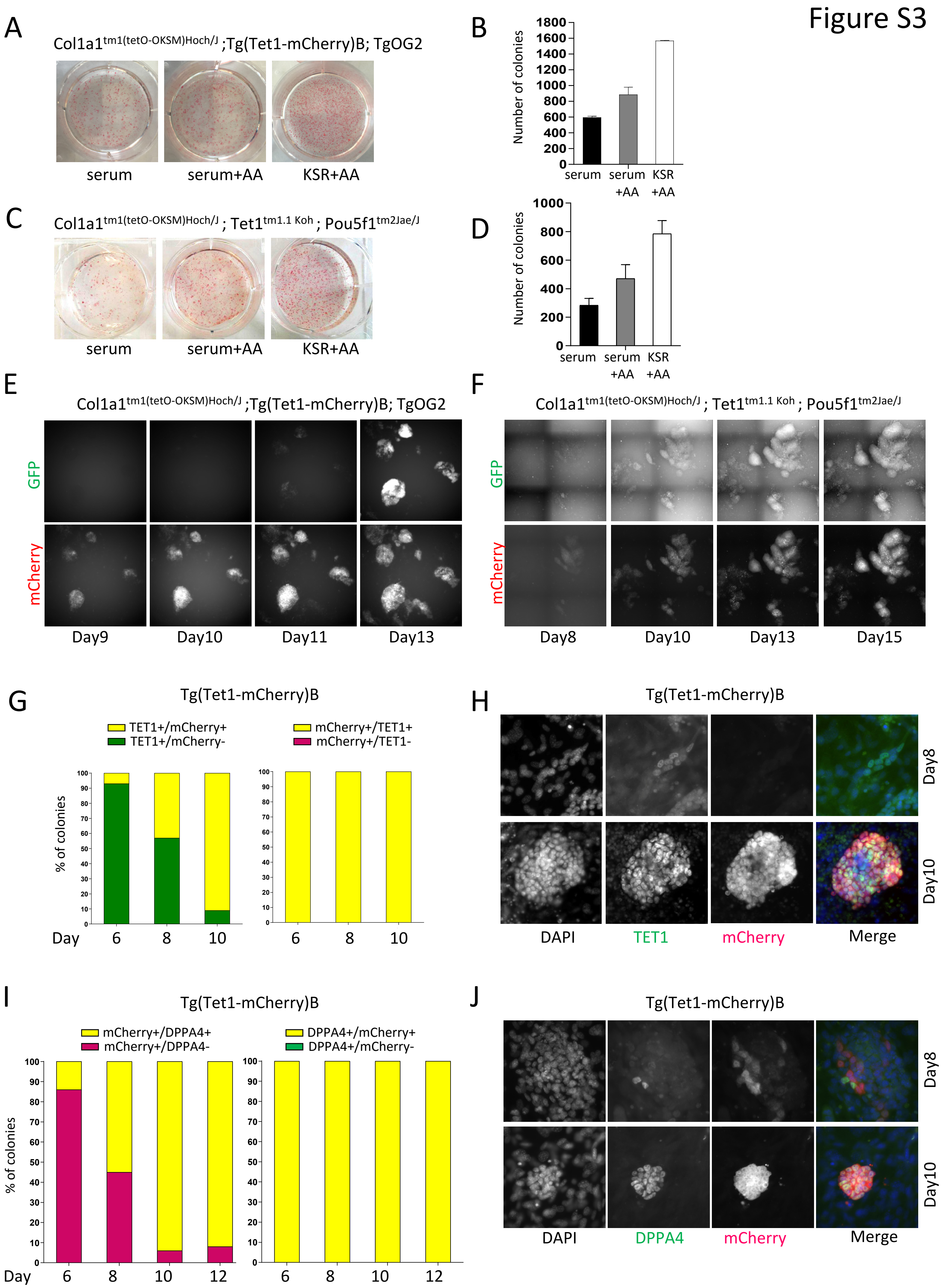
Figure S3
A
Col1a1tm1(tetO-OKSM)Hoch/J ;Tg(Tet1-mCherry)B; TgOG2
serum
serum+AA
KSR+AA
B
Number of colonies
1800
1600
1400
1200
1000
800
600
400
200
0
serum
serum +AA
KSR +AA
C
Col1a1tm1(tetO-OKSM)Hoch/J ; Tet1tm1.1 Koh ; Pou5f1tm2Jae/J
serum
serum+AA
KSR+AA
D
Number of colonies
1000
800
600
400
200
0
serum
serum +AA
KSR +AA
E
Col1a1tm1(tetO-OKSM)Hoch/J ;Tg(Tet1-mCherry)B; TgOG2
GFP
mCherry
Day9
Day10
Day11
Day13
F
Col1a1tm1(tetO-OKSM)Hoch/J ; Tet1tm1.1 Koh ; Pou5f1tm2Jae/J
GFP
mCherry
Day8
Day10
Day13
Day15
G
Tg(Tet1-mCherry)B
TET1+/mCherry+
TET1+/mCherry-
mCherry+/TET1+
mCherry+/TET1-
% of colonies
Day
6
8
10
H
Tg(Tet1-mCherry)B
Day8
Day10
DAPI
TET1
mCherry
Merge
I
Tg(Tet1-mCherry)B
mCherry+/DPPA4+
mCherry+/DPPA4-
DPPA4+/mCherry+
DPPA4+/mCherry-
% of colonies
Day
6
8
10
12
J
Tg(Tet1-mCherry)B
Day8
Day10
DAPI
DPPA4
mCherry
Merge

**Figure S3. Related to Figure 3. Activation dynamics in different reprogramming conditions and relative to NANOG and DPPA4.** (A) Images of alkaline phosphatase (AP)-stained colonies of Tg(*Tet1*-mCherry)B; Tg(*Pou5f1*-EGFP)Mnn after reprogramming at day15 in different media conditions. (B) Colony counts of staining as shown in A (n=2 replicate wells; error bars indicate SD). (C-D) Same as A-B for the *Tet1*$^{\mathrm{tm1.1Koh}}$; *Pou5f1*$^{\mathrm{tm2Jae/J}}$ line. (E-F) Representative fluorescence images of mCherry and GFP reactivation during the time course of reprogramming in KSR+AA media of Tg(*Tet1*-mCherry)B; Tg(*Pou5f1*-EGFP)Mnn (E) and *Tet1*$^{\mathrm{tm1.1Koh}}$; *Pou5f1*$^{\mathrm{tm2Jae/J}}$ dual reporter cells (F). (G-J) Percentage of colonies and representative images for TET1 and mCherry (G-H) and DPPA4 and mCherry (I-J) co-immunofluorescence staining during reprogramming in serum+AA media.

Figure S4

A

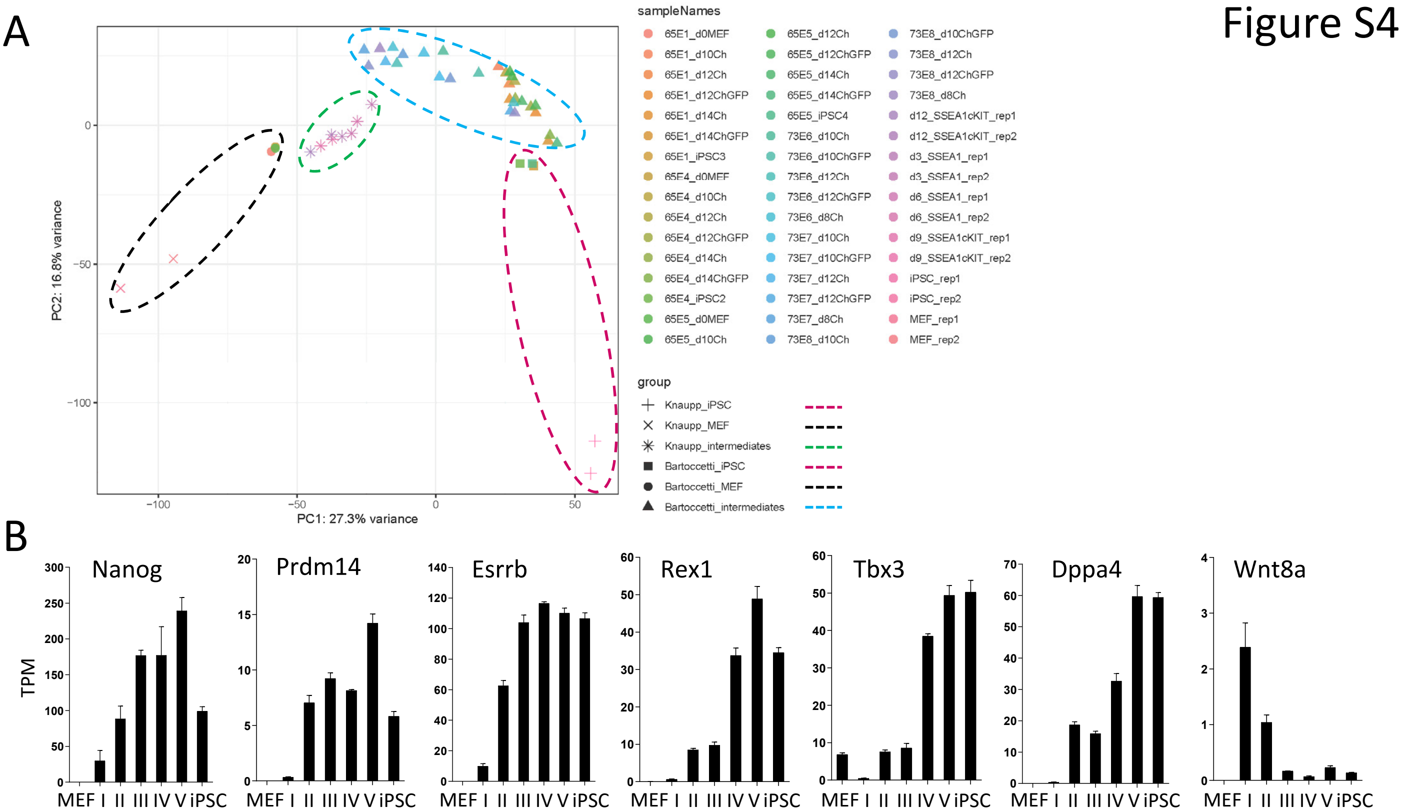

C

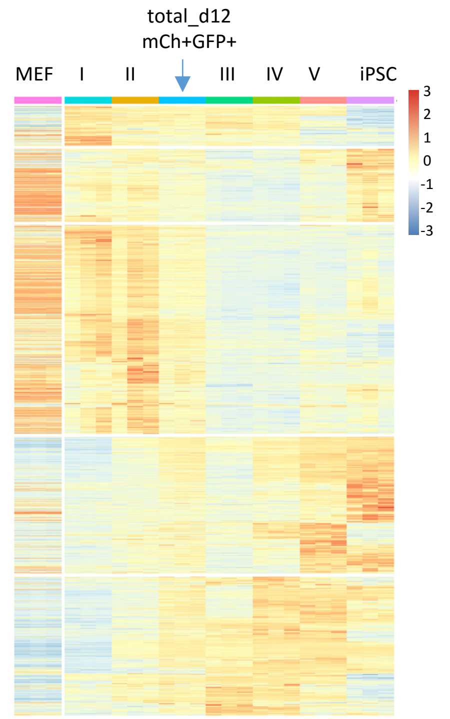

D

GO I - II

| Biological process | Adj p-value |
|---|---|
| pattern specification process | 2.5e-10 |
| regionalization | 3.3e-09 |
| embryonic organ development | 1.3e-08 |
| anterior/posterior pattern specification | 1.5e-08 |
| cell fate commitment | 5.4e-08 |

GO II - III

| Biological process | Adj p-value |
|---|---|
| angiogenesis | 8.3e-44 |
| positive regulation of cell migration | 5.0e-32 |
| positive regulation of cell motility | 5.5e-32 |
| positive regulation of cellular component movement | 9.2e-32 |
| regulation of vasculature development | 3.6e-27 |

GO III - IV

| Biological process | Adj p-value |
|---|---|
| male meiotic nuclear division | 1.8e-11 |
| meiotic nuclear division | 3.0e-09 |
| meiotic cell cycle process | 1.0e-08 |
| DNA methylation involved in gamete generation | 5.9e-08 |
| male gamete generation | 1.2e-07 |

GO IV - V

| Biological process | Adj p-value |
|---|---|
| positive regulation of cell motility | 9.2e-11 |
| positive regulation of cellular component movement | 9.2e-11 |
| positive regulation of cell migration | 4.1e-10 |
| positive regulation of cell-cell adhesion | 5.7e-09 |
| regulation of system process | 6.2e-08 |

E

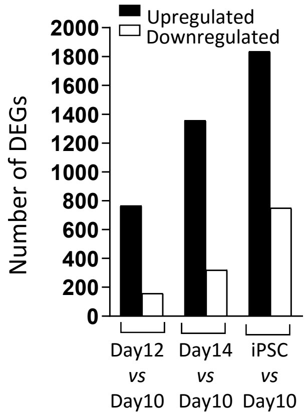

F

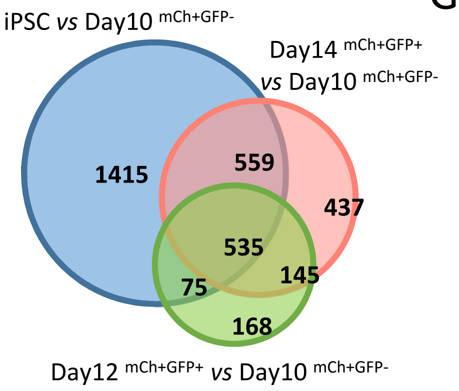

G

| | |
|---|---|
| Taf7l | Tex12 |
| Hsf2bp | Naa11 |
| Fam178b | 1700029P11Rik |
| Sycp2 | Dpep3 |
| Dazl | Ddx4 |
| Slc25a31 | Fkbp6 |
| Ly6k | Hormad1 |
| Sycp3 | Rpl10l |
| Rhox13 | 8030474K03Rik |
| Trim52 | Asz1 |
| Sycp1 | Mael |
| Stk31 | Gm7061 |
| Mov10l1 | 4933416C03Rik |
| D1Pas1 | Hsf5 |
| 1700013H16Rik | |

**Figure S4. Related to Figure 4. Transcriptome changes in reprogramming intermediates sorted based on *Tet1* and *Oct4* reactivation.** (A) Combined gene expression PCA plot of populations identified in this study combined with populations identified by Knaupp et al. Note that our datasets were generated by mRNA-seq whereas Knaupp's datasets were from total RNA-seq, such that the latter dataset occupies a broader space on the combined PCA (see Methods). Nonetheless, the relative positions of starting cells (MEFs), both sets of intermediates and terminal iPSCs can be discerned. (B) Gene expression levels (TPM) of pluripotency-related genes. Values are shown as mean ± SEM (n=3 biological replicates). (C) Heat map of DEGs already shown in Figure 4C with "totalx2" d12 mCh+GFP+ samples included. (D) Top 5 gene ontology (GO) terms associated with stage transitions as defined in Figure 4F. (E) Total number of DEGs upregulated or downregulated in "naivex2" samples relative to "naivex2" mCh+GFP- fractions at day 10 (n_d10mCh) ($|\log_2 FC|>1.5$). (F) Venn diagram overlap of DEG genes as defined in E. (G) List of 29 overlapping genes from Figure 4G.

### Figure S5

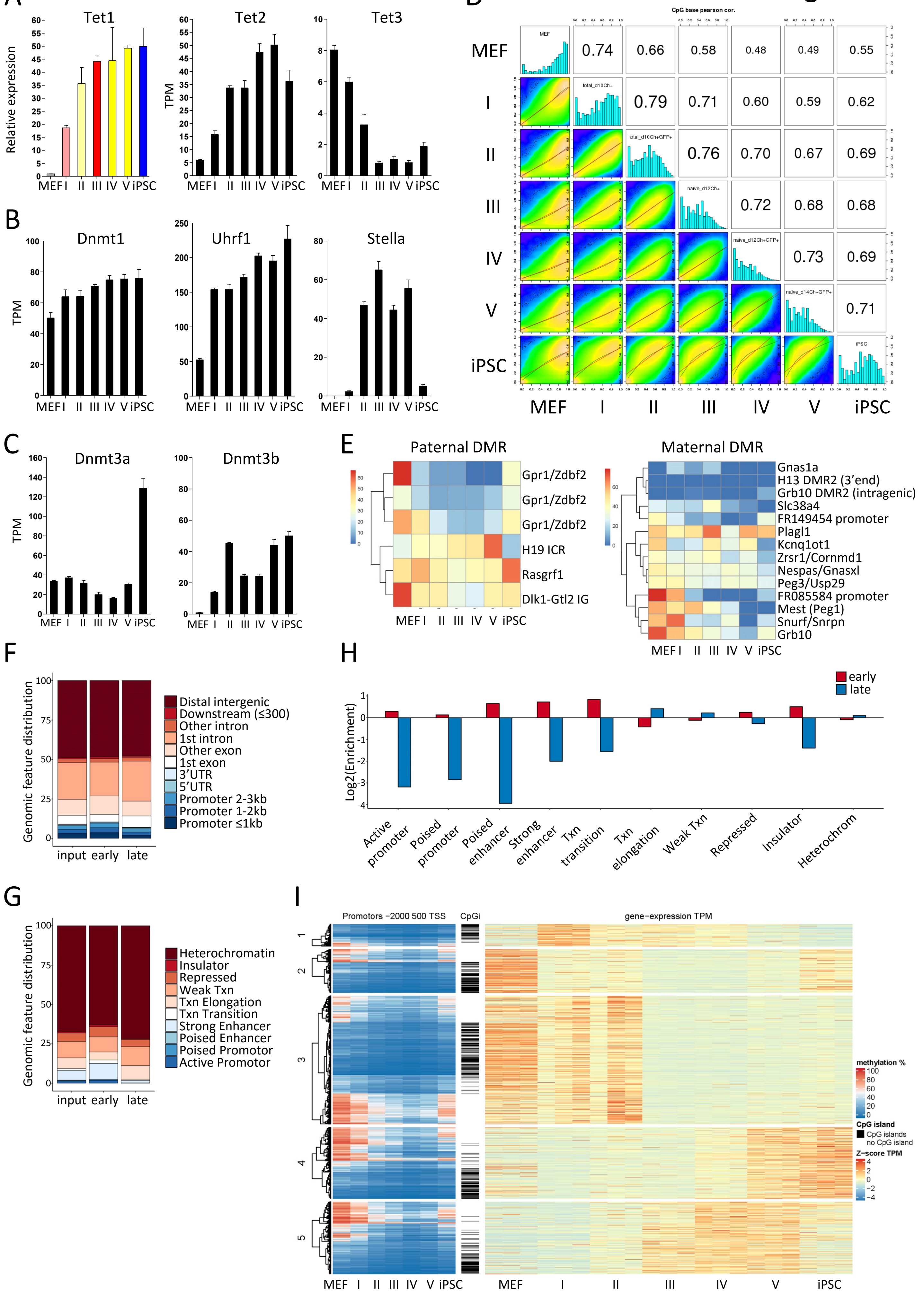

A
Tet1
Relative expression
MEF I II III IV V iPSC
Tet2
TPM
MEF I II III IV V iPSC
Tet3
MEF I II III IV V iPSC
B
Dnmt1
TPM
MEF I II III IV V iPSC
Uhrf1
MEF I II III IV V iPSC
Stella
MEF I II III IV V iPSC
C
Dnmt3a
TPM
MEF I II III IV V iPSC
Dnmt3b
MEF I II III IV V iPSC
D
CpG base pearson cor.
MEF
I
II
III
IV
V
iPSC
MEF
total_d10Ch+
total_d10Ch+GFP+
naive_d12Ch+
naive_d12Ch+GFP+
naive_d14Ch+GFP+
iPSC
0.74
0.66
0.58
0.48
0.49
0.55
0.79
0.71
0.60
0.59
0.62
0.76
0.70
0.67
0.69
0.72
0.68
0.68
0.73
0.69
0.71
MEF I II III IV V iPSC
E
Paternal DMR
Gpr1/Zdbf2
Gpr1/Zdbf2
Gpr1/Zdbf2
H19 ICR
Rasgrf1
Dlk1-Gtl2 IG
MEF I II III IV V iPSC
Maternal DMR
Gnas1a
H13 DMR2 (3'end)
Grb10 DMR2 (intragenic)
Slc38a4
FR149454 promoter
Plagl1
Kcnq1ot1
Zrsr1/Cornmd1
Nespas/Gnasxl
Peg3/Usp29
FR085584 promoter
Mest (Peg1)
Snurf/Snrpn
Grb10
MEF I II III IV V iPSC
F
Genomic feature distribution
Distal intergenic
Downstream (≤300)
Other intron
1st intron
Other exon
1st exon
3'UTR
5'UTR
Promoter 2-3kb
Promoter 1-2kb
Promoter ≤1kb
input early late
H
early
late
Log2(Enrichment)
Active promoter
Poised promoter
Poised enhancer
Strong enhancer
Txn transition
Txn elongation
Weak Txn
Repressed
Insulator
Heterochrom
G
Genomic feature distribution
Heterochromatin
Insulator
Repressed
Weak Txn
Txn Elongation
Txn Transition
Strong Enhancer
Poised Enhancer
Poised Promotor
Active Promotor
input early late
I
Promotors −2000 500 TSS
CpGi
gene−expression TPM
1
2
3
4
5
MEF I II III IV V iPSC
MEF I II III IV V iPSC
methylation %
CpG island
CpG islands
no CpG island
Z−score TPM

**Figure S5. Related to Figure 5. Methylome changes in reprogramming intermediates.** (A-C) Gene expression levels as relative expression by Q-PCR (*Tet1*) or TPM values (all the other genes) at intermediate stages, MEFs and iPSCs. Expression values are means ± SEM (n=3 biological replicates). (D) Pearson correlation of CpG methylation levels among reprogramming intermediates, MEF and iPSC. Histograms show distribution of CpGs classified by methylation levels in 5% bins. (E) Heat map clusters showing methylation levels of maternal and paternal imprint control regions or DMRs (base coverage $\geq$ 5). (F) Genomic features distribution associated with Figure 5H. (G-H) Distribution (G) and enrichment (H) of ESC specific functional annotation of 1-kb tiles associated with early and late changes in CpG methylation as compared to all 1-kbtiles from Figure 5G. (I) CpG methylation levels across promoter regions (-2kb to +0.5 kb of TSS) of genes within DEG clusters shown in Figure 4C (left) plotted against heat map of TPM values (right). Only promoter regions that fulfilled coverage thresholds for methylation calculations are shown. Each row represents a gene ordered by its position in the promoter CpG methylation cluster map.

### Figure S6

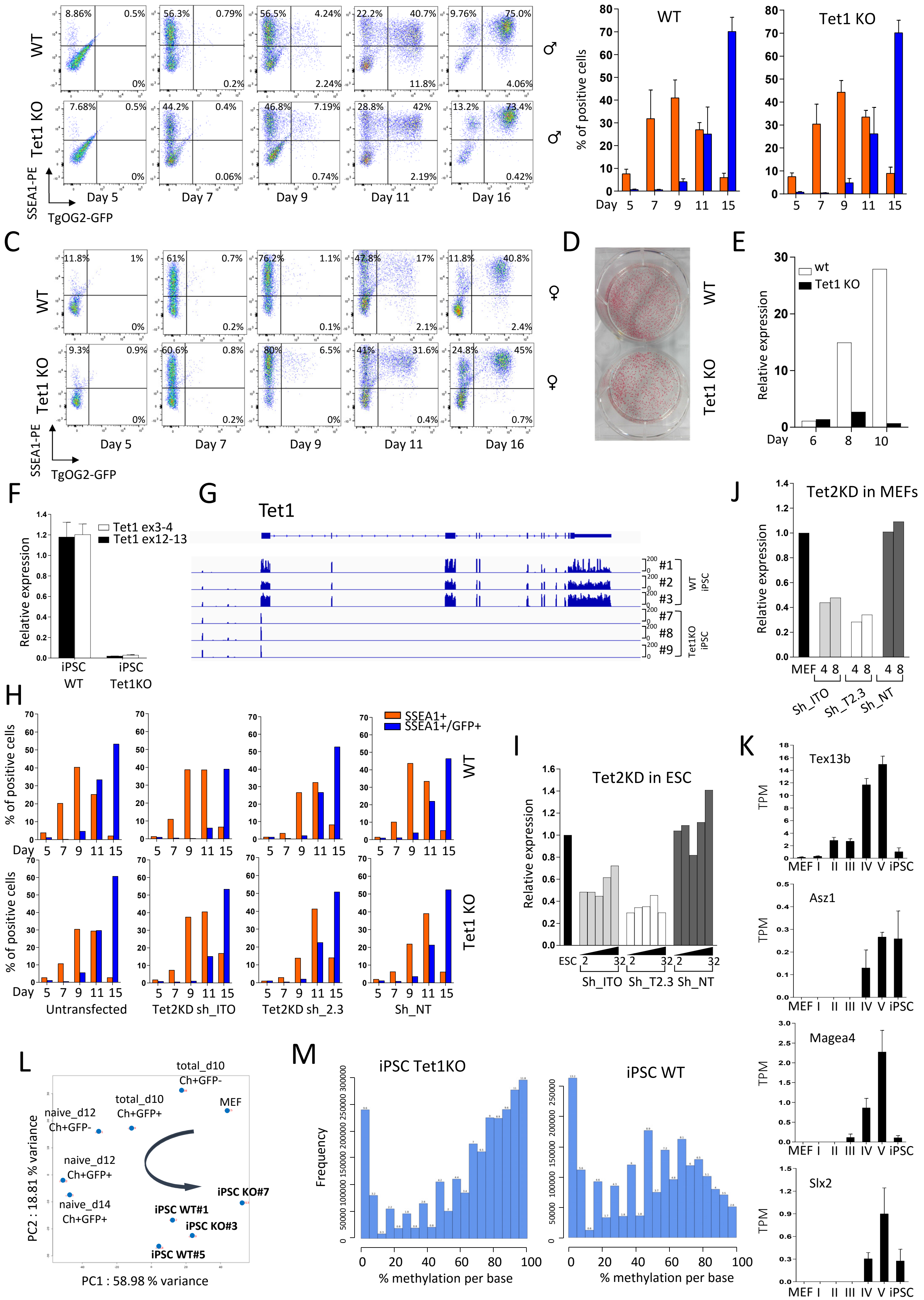

A
Tet1tm1Koh x Tg(Pou5f1-EGFP)
WT
Tet1 KO
SSEA1-PE
TgOG2-GFP
Day 5
Day 7
Day 9
Day 11
Day 16
B
SSEA1+
SSEA1+/GFP+
WT
Tet1 KO
% of positive cells
C
D
E
wt
Tet1 KO
Relative expression
F
Tet1 ex3-4
Tet1 ex12-13
iPSC WT
iPSC Tet1KO
G
Tet1
J
Tet2KD in MEFs
MEF
Sh_ITO
Sh_T2.3
Sh_NT
H
Untransfected
Tet2KD sh_ITO
Tet2KD sh_2.3
Sh_NT
I
Tet2KD in ESC
ESC
K
Tex13b
Asz1
Magea4
Slx2
TPM
L
total_d10 Ch+GFP-
total_d10 Ch+GFP+
MEF
naive_d12 Ch+GFP-
naive_d12 Ch+GFP+
naive_d14 Ch+GFP+
iPSC KO#7
iPSC WT#1
iPSC KO#3
iPSC WT#5
PC1 : 58.98 % variance
PC2 : 18.81 % variance
M
iPSC Tet1KO
iPSC WT
Frequency
% methylation per base

**Figure S6. Related to Figure 6. Reprogramming in the absence of Tet1.** (A) Representative flow cytometry analysis of *Tet1* KO (*Tet1*$^{tm1Koh}$) and wt littermate cell lines harboring TgOG2, both males, harvested at different time points during reprogramming. Cells were reprogrammed in KSR+AA media and stained with SSEA1 antibody. (B) Percentages of gated cell fractions in replicate experiments of A are shown as mean ± SEM (n=3 independent littermate pairs of wt and KO). (C) Same as A with female cells. (D) AP staining of the experiment shown in C at day15 of reprogramming. (E) Relative expression of *Tet1* in wt and *Tet1*-KO at different time points during reprogramming (primers for *Tet1* detect exons 12-13). (F) Relative expression of *Tet1* in wt and KO iPSCs measured by qPCR. Primers amplify coding regions in exons 3-4 and 12-13 of *Tet1*. Expression values are means ± SEM (wt n=3, *Tet1*-KO n=6). (G) IGV RNA-seq tracks over the *Tet1* gene in 3 wt and 3 KO iPSC clonal lines. (H) Percentages of gated cell fractions of wt and *Tet1* KO cell lines containing TgOG2 after transfection with two different Tet2sh_RNA, non-targeting (NT)_shRNA or untransfected at different time points during reprogramming. Cells were stained with SSEA1 antibody. (I-J) Relative expression of *Tet2* in ESCs (I) and MEFs (J) transfected with two different Tet2_shRNA. Sh_NT was used as control. Values on the x- axes indicate µl of virus supernatant used in 2-fold increments. (K) Gene expression levels (TPM) of representative genes identified in Figure 6D. Values are shown as mean ± SEM (n=3). The plot for *Tex12* is shown in Figure S7A. (L) PCA based on single CpG methylation levels of 5 intermediates together with 2 wt and 2 *Tet1*-KO iPSC lines. WT#5 and *Tet1*-KO#3 are sequenced at low (˜3-fold) genome coverage. Arrow indicates the direction from MEF to iPSC. (M) Distribution of CpGs classified by methylation levels in *Tet1* KO and wt iPSCs.

### Figure S7

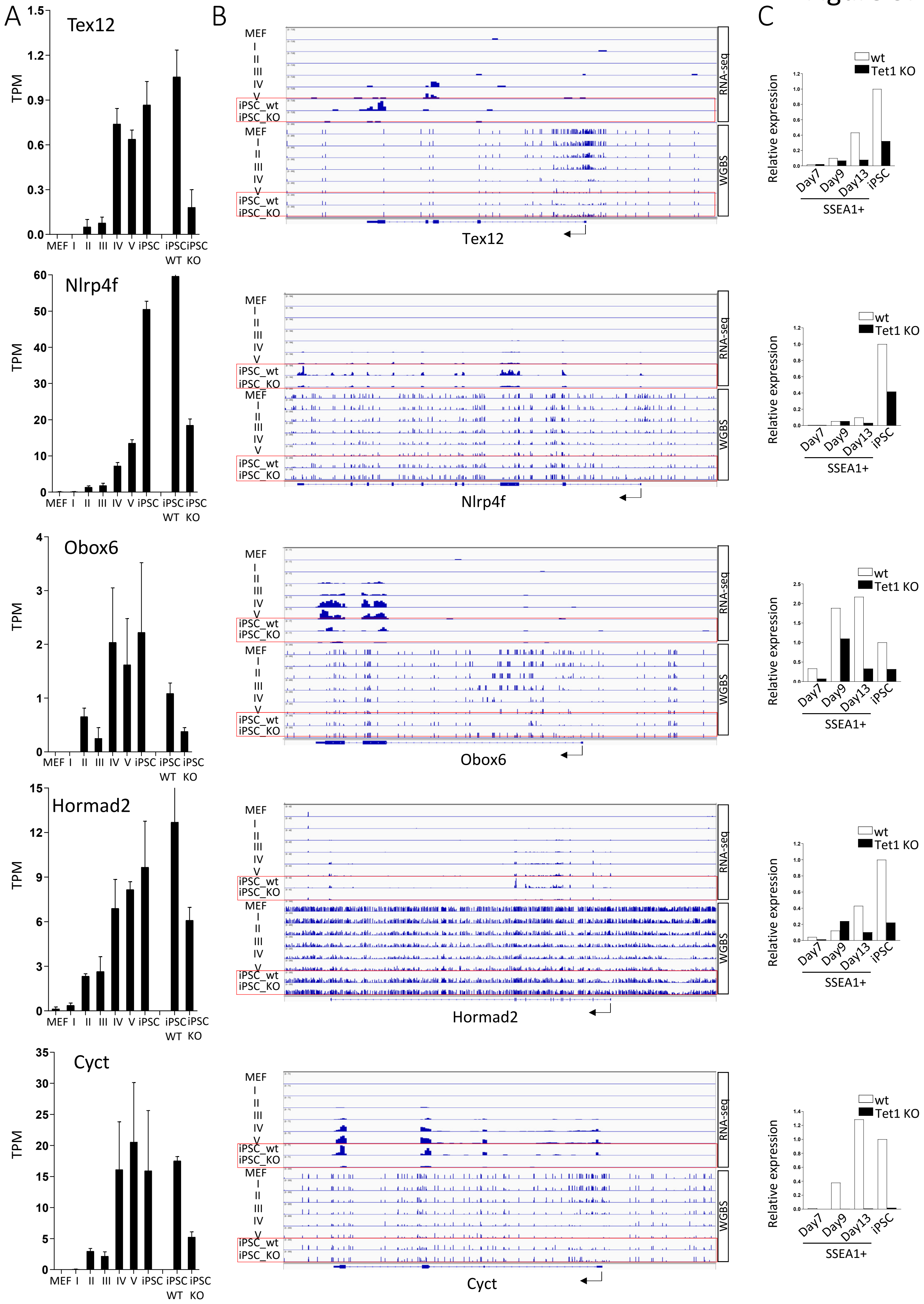
A
B
C
Tex12
Nlrp4f
Obox6
Hormad2
Cyct
TPM
MEF
I
II
III
IV
V
iPSC
iPSC WT
iPSC KO
iPSC_wt
iPSC_KO
RNA-seq
WGBS
wt
Tet1 KO
Relative expression
Day7
Day9
Day13
SSEA1+

**Figure S7. Related to Figure 6. *Tet1* dependent genes.** (A) TPM values for *Tex12, Nlrp4f*, *Obox6*, *Hormad2* and *Cyct* in stage intermediates, MEFs and wt and *Tet1*-KO iPSCs. Expression values are means ± SEM (n=3). (B) IGV RNA-seq and WGBS tracks over these 5 genes in stage intermediates, MEF, wt and *Tet1* KO iPSC. (C) Relative expression measured by Q-PCR in SSEA1+ sorted populations at different time points during reprogramming in KSR+AA.

| **Souther blot primers** | | |
|---|---|---|
| **Name** | **Sequence (5’→3’)** | |
| Probe4_Fw | CTGCTGAGCCTTATCACTATCC | |
| Probe4_Rv | CGAGAGTTGGCAGAATGAAGA | |
| **Genotyping primers** | | |
| **Name** | **Sequence (5’→3’)** | **Mouse strain** |
| Col A1 WT_F | GCACAGCATTGCGGACATGC | B6;129S4-Col1a1tm1(tetO-Pou5f1,-Klf4,-Sox2,-Myc)Hoch/J |
| Col B Rv | CCCTCCATGTGTGACCAAGG | |
| Col C1 Mut_Rv | GCAGAAGCGCGGCCGTCTGG | |
| Tet1P13-3’seqFw | TGGAAGCCTCTTGCATCATC | B6;129S-Tg(Tet1-mCherry)B |
| mCherryseq164Rv | TTGGTCACCTTCAGCTTGG | |
| Tet1c5-Gtype-Rv | GGGTGTCTAAAGACAGCTACA | |
| Pou5f1tm2Jae_wt_F | CAAGGCAAGGGAGGTAGACA | B6;129S4-Pou5f1tm2Jae/J |
| Pou5f1tm2Jae_wt_R | TGCCAGACAATGGCTATGAG | |
| Pou5f1tm2Jae_mutR | CCAAAAGACGGCAATATGGT | |
| EGFP-496F | TCAAGATCCGCCACAACATC | B6;CBA-Tg(Pou5f1-EGFP)2Mnn/J |
| EGFP_R | CGCTTTACTTGTACAGCTCGT | |
| Tet1-1F (Tet1tm1) | TTAGACCCCAAACTCAGGTGAC | B6; Tet1tm1.1Koh and B6; Tet1tm1Koh |
| Tet1-2R (Tet1tm1) | GTTTTCCGGGGTTCACTGCCTT | |
| LacZ-6R (Tet1tm1) | CGGATTGACCGTAATGGGATAG | |
| CreTet1tm1.1-Gtype-5R | AGCTTCAGCCTCTGCTTGATCT | |

| qPCR primers | |
|---|---|
| **Name** | **Sequence (5’→3’)** |
| Gapdh Fw | ACCACAGTCCATGCCATCAC |
| Gapdh Rev | CACCACCCTGTTGCTGTAGCC |
| Tet1 exon 1a Fw | CTGCCTCTTCTACGGGAACATTCG |
| Tet1 exon 1b Fw | TCTGTCCTGGCTGAGTGTCTTCAT |
| Tet1 exon 2 Rev | GCCTGCTTTGATGTCTTCGTCTTC |
| Tet1 Exon12-13 Fw | GAGCCTGTTCCTCGATGTGG |
| Tet1 Exon12-13 Rev | CAAACCCACCTGAGGCTGTT |
| Tet1 Exon3-4 Fw | GAGGGAAAAGAAGCCCAAAG |
| Tet1 Exon3-4 Rev | CGCCTGCATTCTTCCTTACA |
| Tet1 Fw | GAGCCTGTTCCTCGATGTGG |
| Tet1 Rev | CAAACCCACCTGAGGCTGTT |
| Tet2 Fw | AACCTGGCTACTGTCATTGCTCCA |
| Tet2 Rev | ATGTTCTGCTGGTCTCTGTGGGAA |
| Tet3 Fw | TTCAACGGCTGCAAATATGCTCGG |
| Tet3 Rev | ACTACTGACCTTGGCGTTCTGGTT |
| Dnmt1 Fw | CCTAGTTCCGTGGCTACGAGGAGAA |
| Dnmt1 Rev | TCTCTCTCCTCTGCAGCCGACTCA |
| Dnmt3a Fw | GCCGAATTGTGTCTTGGTGGATGACA |
| Dnmt3a Rev | CCTGGTGGAATGCACTGCAGAAGGA |
| Dnmt3b Fw | GCCCATGCAATGATCTCTCT |
| Dnmt3b Rev | CCAGAAGAATGGACGGTTGT |
| Blimp1 Fw | AGCATGACCTGACATTGACACC |
| Blimp1 Rev | CTCAACACTCTCATGTAAGAGGC |
| Stella Fw | AGGCTCGAAGGAAATGAGTTTG |
| Stella Rev | TCCTAATTCTTCCCGATTTTCG |
| Prdm14 Fw | GCATATACCCTACCCGCTTTC |
| Prdm14 Rev | CAAACGGATTGGAGGTTGAT |

| | |
|---|---|
| Rex1 Fw | CAGCTCTGCACACAGAAGA |
| Rex1 Rev | ACTGATCCGCAAACACCT |
| Tbx3 Fw | TGGAACCCGAAGAAGACGTAG |
| Tbx3 Rev | TACCCCGCTTGTGAAACTGG |
| Klf4 Fw | CCAGCAAGTCAGCTTGTGAA |
| Klf4 Rev | GGGCATGTTCAAGTTGGATT |
| Esrrb Fw | TTTCTGGAACCCATGGAGAG |
| Esrrb Rv | AGCCAGCACCTCCTTCTACA |
| Fgf5 Fw | AAAGTCAATGGCTCCCACGAA |
| Fgf5 Rev | CTTCAGTCTGTACTTCACTGG |
| Otx2 Fw | CCACTTCGGGTATGGACTTG |
| Otx2 Rev | GTCCTCTCCCTTCGCTGTTT |
| Wnt8a Fw | CATGTACGCAGTCACCAAGAA |
| Wnt8a Rev | CATCCTTCCCTTTCTCCAAAC |
| Tex12 Fw | GTGCCTATGCCAGCTAAAAGT |
| Tex12 Rv | TATAATGTGCCAAATATTTGACCCTC |
| Nlrp4f Fw | GCCCATGGTTCTTGCATCA |
| Nlrp4f Rv | ATCAGAGGCAGTGTGGAAGCA |
| Obox6 Fw | CACAGCAAATGAGATCCAGAT |
| Obox6 Rv | ATACCTGGCACTATCACAGGC |
| Hormad2 Fw | CTACTGAGATAGCTCATCAGGG |
| Hormad2 Rv | GACTGTCACTGGTTCGCTGACC |
| Cyct Fw | GCACAGCAGTTGCACCTTAC |
| Cyct Rv | TGTATTCACAATCACAGCAGCC |
| **Bisulfite primers** | |
| **Name** | **Sequence (5’→3’)** |
| Oct4-Pr-bis_F1 | GAGGATTGGAGGTGTAATGGTTGTT |
| Oct4-Pr-bis_R1 | CTACTAACCCATCACCCCCACCTA |
| Oct4-DE-bis_F1 | GAATAGAATTTTAAAAGGGTTTTTTG |
| Oct4-DE-bis_Fnest | TTGAGTTTTTTTTTAGTTTTTAGTTTTG |

| | |
|---|---|
| Oct4-DE-bis_R1 | TAATTCAAAAATTCCCTACATAACCT |
| Oct4-DE-bis_Rnest | ACTAAAATTAAAAATATATACCACTCTACC |
| Tet1 D2-Bis_F | GGGAAAAAGTGGATTATATTTAGTTTAG |
| Tet1 D2-Bis_R | CATAAACACACACATACAAAAAAAA |
